# Genomic neighbor typing for bacterial outbreak surveillance

**DOI:** 10.1101/2022.02.05.479210

**Authors:** Eike Steinig, Miranda Pitt, Izzard Aglua, Annika Suttie, Andrew Greenhill, Christopher Heather, Cadhla Firth, Simon Smith, William Pomat, Paul Horwood, Emma McBryde, Lachlan Coin

## Abstract

Genomic neighbor typing enables heuristic inference of bacterial lineages and phenotypes from nanopore sequencing data. However, small reference databases may not be sufficiently representative of the diversity of lineages and genotypes present in a collection of isolates. In this study, we explore the use of genomic neighbor typing for surveillance of community-associated *Staphylococcus aureus* outbreaks in Papua New Guinea (PNG) and Far North Queensland, Australia (FNQ). We developed `Sketchy`, an implementation of genomic neighbor typing that queries exhaustive whole genome reference databases using MinHash. Evaluations were conducted using nanopore read simulations and six species-wide reference sketches (4832 - 47616 genomes), as well as two *S. aureus* outbreak data sets sequenced at low depth using a sequential multiplex library protocol on the MinION (n = 160, with matching Illumina data). Heuristic inference of lineages and antimicrobial resistance profiles allowed us to conduct multiplex genotyping *in situ* at the Papua New Guinea Institute of Medical Research in Goroka, on low-throughput Flongle adapters and using multiple successive libraries on the same MinION flow cell (n = 24 - 48). Comparison to phylogenetically informed genomic neighbor typing with `RASE` on the dominant outbreak sequence type suggests slightly better performance at predicting lineage-scale genotypes using large sketch sizes, but inferior performance in resolving clade-specific genotypes (methicillin resistance). `Sketchy` can be used for large-scale bacterial outbreak surveillance and in challenging sequencing scenarios, but improvements to clade-specific genotype inference are needed for diagnostic applications. `Sketchy` is available open-source at: https://github.com/esteinig/sketchy

---

**E**pidemiological and clinical features of infectious diseases, such as strain provenance and antimicrobial susceptibility, are valuable targets for decision makers, but their timely inference from genomic data is challenging. Fast methods for genotyping are especially relevant for bacterial pathogens for which faithful genome assembly requires reasonable genome coverage. However, whole genomes often cannot be assembled easily from complex metagenomic samples, including blood and lower respiratory infections where the causative pathogen may be at low abundance.

Nanopore sequencing is particularly suited for rapid characterisation of pathogen genomes, with potential to be conducted on-site rather than sent to a reference lab, as reads can be streamed from the device and analysed on mobile computing platforms (1–4). Several methods for bacterial pathogens characterisation on nanopore platforms has been developed over the past few years, including pipelines for batch assembly and marker detection (5, 6), novel algorithms for streaming assembly and genotyping (7–9), and sensitive approaches to antimicrobial resistance prediction, as well as taxonomic identification (10–12). Studies that assessed clinical specimens have focused on samples with low abundance of host nucleic acids, high bacterial loads, and those in which nanopore sequencing was supported with short-read sequencing (6, 13–15). Strain-level genotyping from lower respiratory infections and cystic fibrosis patients was particularly efficient when preceded by host nucleic acid depletion(13) or enriched by culture (14).

In pursuit of rapid genotype inference, Břinda *et al*. developed a heuristic principle termed “genomic neighbor typing” (16). Antibiotic resistance phenotypes (minimum inhibitory concentrations) and lineage membership could be inferred using *k*-mer matching against a database of whole genome sequences, including their phylogenetic relationships `RASE`). Using genomic neighbor typing, heuristic inference of genome-associated traits was possible within minutes of starting sequencing. Genomic neighbor typing could thus be used for massively parallel genotyping, requiring only standard nucleic acid extraction and multiplex library protocols to survey lineage and genotype composition of an bacterial outbreak, where complete genomes may be difficult to produce at scale.

A critical component of genomic neighbor typing is sufficient representation of genome diversity in the reference database. Břinda *et al*. constructed reference sets from local and national collections to demonstrate the principle using *S. pneumoniae* (n = 616) and *Neisseria gonorrhoeae* (n = 1102). However, for clinical applications and in particular for outbreak scenarios, a small reference database of the globally available sequence space for a pathogen may be insufficiently representative of species-wide diversity and miss important lineages or sublineage genotypes that may have entered local epidemiological space. In one such outbreak scenario in Papua New Guinea, community-associated MRSA infections in Kundiawa (Simbu Province) and Goroka (Eastern Highlands Province) had been tracked over multiple years, but lineage provenance and genotype identity had been unknown (17).

No sequencing data on *S. aureus* from Papua New Guinea exists, so that the application of genomic neighbor typing to survey the lineage and genotype composition of the outbreak would require a sufficiently representative lineage database to account for the absence of prior data from the region.

In addition to the outbreak in the remote highland provinces of Papua New Guinea, we had become aware of escalating *S. aureus* infections in remote communities of Far North Queensland, which borders Papua New Guinea through the Torres Strait Islands (18, 19). We collected a snap-shot of strains from Far North Queensland communities (Cape York Peninsula, Cairns and Hinterland, Torres Strait Island) in 2019 (20). This presents a realistic scenario, where surveying cross-border bacterial outbreaks using a comprehensive genomic neighbor typing approach on nanopore devices could provide lineage and genotype data important for deciphering geographical provenance (lineage attribution) and antibiotic susceptibilities of dominant outbreak lineages, potentially informing treatment options. Furthermore, we were interested in demonstrating that genomic neighbor typing is working under realistic sequencing conditions, including on site in Goroka (Eastern Highlands Province) where access to sequencing infrastructure is not available. In addition, we wanted to assess using genomic neighbor typing as a cost-effective approach, for example by using successive library sequencing protocols or Flongle.

Large reference databases may be required to ensure a ‘hypothesis-agnostic’ genomic neighbor typing approach. However, maintaining a relatively small resource profile while using large reference databases of bacterial whole genome sequences requires an approximate database construction and read matching approach that can accommodate tens to potentially hundreds of thousands of genomes. MinHash, a variant of locality-sensitive hashing originally used for detection of near-duplicate websites or images (21), has been extensively used in genomics since its implementation in `Mash` (22, 23). Computing min-wise shared hashes between reference and query sketches (23, 24) presents a simple method to implement genomic neighbor typing with comprehensive lineage and genotype representation and without the need for phylogenetic trees as required by `RASE`. In addition, an implementation of genomic neighbor typing that uses genotypes, instead of culture-based phenotypes (such as MIC values) would allow for the construction of reference sketches entirely from public genome collections.

In this study, we evaluate genomic neighbor typing with species-wide bacterial pathogen sketches using MinHash. We developed a simple genomic neighbor typing approach using ranked shared hashes and lineage-resolved (‘hypothesis-agnostic’) databases which span the known genomic diversity of a bacterial species and are constructed from public sources (Fig. 1). Our primary aim was to infer lineage and sublineage genotypes from as few reads as possible, and to evaluate the approach on independent outbreak data from remote northern Australia and Papua New Guinea (n = 160, with matching Illumina reference data). We reasoned that genomic neighbor typing could be used for scaling outbreak surveillance through heuristic genotype inference.

**Fig. 1.**
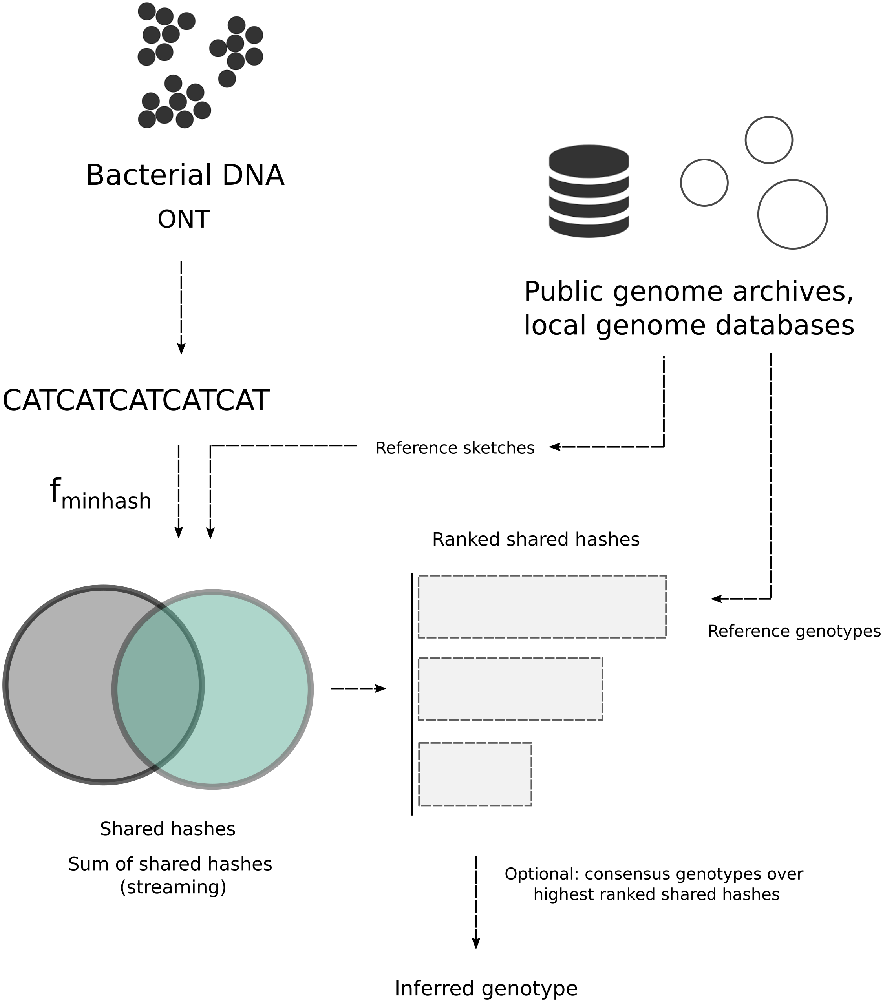
`Sketchy` components: bacterial sequence reads are matched against a reference database constructed from bacterial whole genomes (including from public archives). Shared hashes (or the sum of shared hashes for streaming operations) are then ranked and the genotype of the highest ranking genome match selected for prediction. Alternatively, consensus calls for each genotype feature (binary or multi-label, e.g. trimethoprim resistance or *SCCmec* subtype) can be made over the highest ranking genomes in the reference sketch.

## Results

### Species cross-validation simulation

Whole genome sequences for six species with varying levels of representation in the European Nucleotide Archive (ENA) were collected for reference sketch construction (Table 1) (25). After filtering of assemblies for contamination, completeness and strain heterogeneity, we constructed default (*k* = 16, *s* = 1000) and high resolution (*k* = 16, *s* = 10000) reference sketches for evaluation (Table 1). Sketch databases contained between 4,832 (*Pseudomonas aeruginosa*) and 47,616 genomes (*S. pneumoniae*). Low resolution sketch files were considerably smaller and consumed less memory than their high resolution equivalents (Table 1). We used multi-locus sequence types (MLST) as a proxy for genotype predictions, as MLST data were readily available for all species (25) and representative of the ability to match genomes in the correct genomic neighborhood (lineage) of the reference database. We conducted a cross-validation simulation, for which we sampled 10 genomes (without replacement) from the reference collection of each species across 20 replicates (n = 200) and used reference sketches which did (DB+) or did not (DB-) contain the sampled genomes (Fig. 2, Table 1). Since our primary aim was to call genotypes from as few reads as possible, we evaluated performance (mean proportion correctly classified) at a threshold of 1000 reads (Methods). We assessed two other methods at this threshold for comparison with `Sketchy`: a *k*-mer based MLST allele typer for long reads (`Krocus`) and whole genome assembly with `Flye`, with added polish by `Medaka`.

**Fig. 2.**
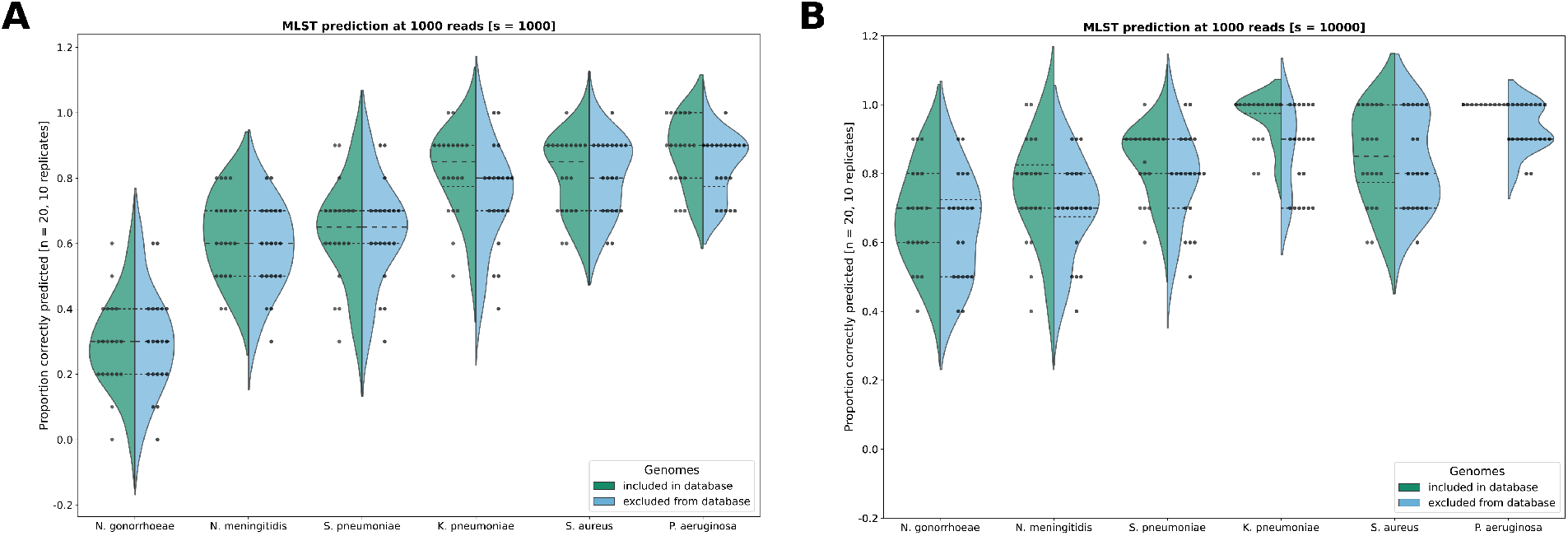
Cross-validation MLST classification success (proportion of correct sequence types, at 1000 reads) for (**A**) default resolution (*s* = 1000) sketches and (**B)** high-resolution (*s* = 10000) sketches across species (violins, data points) showing one half as reference sketches containing the hold-out samples (green, left, DB+) and the other as reference sketches that did not contain the hold-out samples (blue, right, DB-). Cross-validation sampling was conducted using 20 replicates of 10 random samples from the complete reference genome collection and reference sketches (*k* = 16) were built with and without the sampled genomes (Table 1).

**Table 1.** Cross-validation of species sketches using simulated nanopore reads (1000 reads, k = 16, 20 replicates, n = 10)

| Species | n | s | Size (MB) | LZMA (MB) | Time (s) | MEM (GB) | DB+ (%) | DB- (%) |
| --- | --- | --- | --- | --- | --- | --- | --- | --- |
| *Streptococcus pneumoniae* | 47616 | 1000 | 552 | 46 | 5.06 ($\pm 0.16$) | 5.97 ($\pm 0.001$) | 63.5 ($\pm 15.3$) | 63.5 ($\pm 15.3$) |
| *Staphylococcus aureus* | 42461 | 1000 | 492 | 38 | 3.72 ($\pm 0.07$) | 5.32 ($\pm 0.001$) | 79.5 ($\pm 9.44$) | 80.5 ($\pm 11.5$) |
| *Neisseria meningitidis* | 16198 | 1000 | 188 | 16 | 1.48 ($\pm 0.01$) | 2.03 ($\pm 0.001$) | 62.5 ($\pm 12.9$) | 58.5 ($\pm 13.4$) |
| *Klebsiella pneumoniae* | 10072 | 1000 | 117 | 10 | 1.10 ($\pm 0.01$) | 1.26 ($\pm 0.001$) | 82.0 ($\pm 12.8$) | 74.5 ($\pm 15.4$) |
| *Neisseria gonorrhoeae* | 8413 | 1000 | 98 | 7 | 0.81 ($\pm 0.01$) | 1.05 ($\pm 0.001$) | 29.0 ($\pm 13.8$) | 29.5 ($\pm 15.4$) |
| *Pseudomonas aeruginosa* | 4832 | 1000 | 56 | 5 | 0.67 ($\pm 0.01$) | 0.61 ($\pm 0.001$) | 88.0 ($\pm 10.6$) | 83.5 ($\pm 9.33$) |
| *Streptococcus pneumoniae* | 47616 | 10000 | 5720 | 791 | 44.33 ($\pm 3.86$) | 58.82 ($\pm 0.001$) | 83.5 ($\pm 10.5$) | 78.5 ($\pm 13.4$) |
| *Staphylococcus aureus* | 42461 | 10000 | 5101 | 671 | 34.85 ($\pm 1.77$) | 53.34 ($\pm 0.001$) | 84.5 ($\pm 13.6$) | 83.0 ($\pm 13.4$) |
| *Neisseria meningitidis* | 16198 | 10000 | 1945 | 221 | 14.59 ($\pm 0.25$) | 20.35 ($\pm 0.001$) | 75.5 ($\pm 15.4$) | 70.0 ($\pm 14.1$) |
| *Klebsiella pneumoniae* | 10072 | 10000 | 1210 | 139 | 9.33 ($\pm 0.34$) | 12.65 ($\pm 0.001$) | 96.5 ($\pm 6.71$) | 86.0 ($\pm 12.3$) |
| *Neisseria gonorrhoeae* | 8413 | 10000 | 1011 | 112 | 7.98 ($\pm 0.66$) | 10.57 ($\pm 0.001$) | 67.5 ($\pm 14.5$) | 64.5 ($\pm 15.4$) |
| *Pseudomonas aeruginosa* | 4832 | 10000 | 554 | 75 | 4.31 ($\pm 0.02$) | 6.07 ($\pm 0.001$) | 100.0 ($\pm 0.0$) | 93.0 ($\pm 6.57$) |

LZMA (compression), DB+ (random sample retained in reference sketch), DB- (random sample excluded from reference sketch)

Performance was dependent on species, with three out of six species (*N. gonorrhoeae*, *N. meningiditis* and *S. pneumoniae*) showing inferior lineage predictions in the cross-validation assessment, including extremely low performance by *N. gonorrhoeae* (Table 1, Fig. 2). In contrast, *K. pneumoniae*, *S. aureus* and *P. aeruginosa* recovered MLST reasonably well, ranging from 79.5% - 100% (DB+) and 74.5% - 93.0% accuracy (Table 1, Fig. 2). For these species, predictions improved when sampled genomes were contained in the reference sketch (green, DB+) but this trend was not as pronounced for under-performing species (Fig. 1). In all species, predictions improved, often considerably, using higher resolution sketches (*s* = 10000), including accurate predictions using sketches which did not contain the sampled genomes for *P. aeruginosa* (93% ± 6.57) and *K. pneumoniae* (86% ± 12.3) (Table 1).

In species where sketches were able to sufficiently recover lineage, performance stabilised around 200 - 500 reads suggesting that fewer reads than the threshold may be sufficient for genotyping (Fig. S1A). Reference sketch size, memory consumption and prediction times scaled approximately linearly with the number of genomes in the reference sketch and sketch size (Table 1). Predictions at an additional threshold of 10000 reads using the low-resolution sketches (*s* = 1000) showed minor improvements in performance across species (Fig. S1B), indicating that most of the observed error was due to the resolution of the sketching approach, and only some due to the low read threshold chosen for typing. Finally we compared `Sketchy` to allele based *k*-mer matching with `Krocus` and lineage typing from assemblies generated with `Flye`, optionally polished with `Medaka. Krocus` was unable to infer MLST from 1000 reads in all cases (Fig1. S1C). Using assembled genomes, allele typing led to incorrect multi-locus sequence types in all cases, except in a single *P. aeruginosa* assembly with `Flye` (Fig. S1C).

### Genotype surveillance of community-associated outbreaks

We next evaluated Sketchy genotyping on two *S. aureus* outbreaks from remote communities in Papua New Guinea and Far North Queensland (n = 160), which had been sequenced at low-coverage using a dual-library protocol with interspersed nuclease washes (24 strains per MinION flow cell, on a total of 8 flow cells) (4) (Fig. 3, Methods). While most isolates belonged to the Australian ST93-MRSA-IV clone (26, 27) (n = 120), multiple other sequence types were recovered (ST1, ST5, ST15, ST25, ST30, ST45, ST81, ST121, ST243, ST762, n = 35), including several novel sequence types (n = 5) of which some derived from the ST93 outbreak lineage (n = 3). In addition, a version of the *S. aureus* reference sketch was created using a collection of genomes for which we had previously computed antimicrobial resistance calls with `Mykrobe` (n = 34,583), as well as *SCCmec* type and presence of the Panton-Valentine leukocidin locus (PVL). As the outbreak isolates were the first *S. aureus* genomes recovered from Papua New Guinea, these data comprised an independent validation dataset for performance evaluation of the *S. aureus* reference sketch, in the absence of local or regional genome collections.

**Fig. 3.**
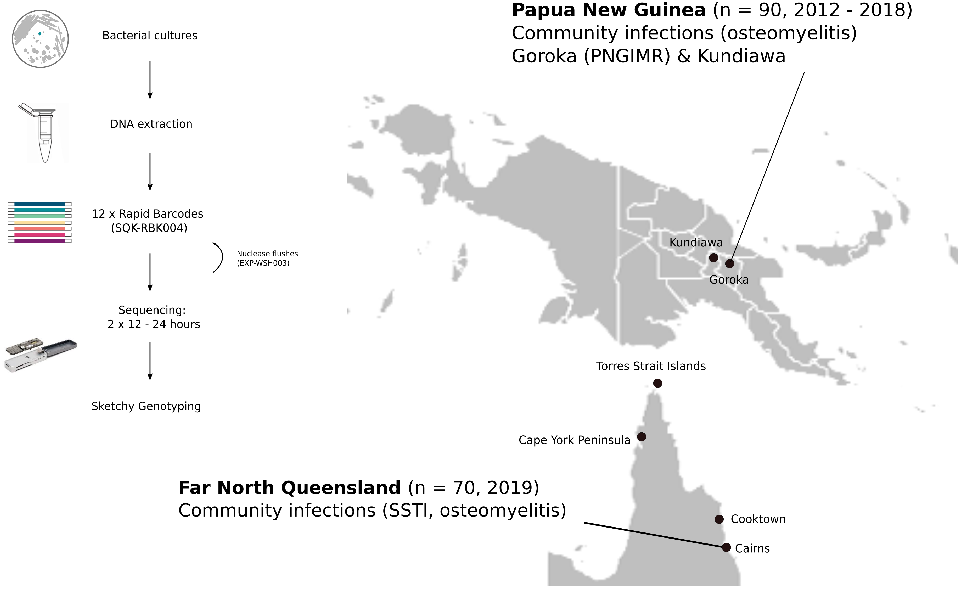
Nanopore validation dataset with matching Illumina data sequenced from community-associated *S. aureus* outbreaks in Papua New Guinea (PNG) and Far North Queensland (FNQ). Sampling of osteomyelitis and skin-and soft tissue infections (SSTI) was conducted in Goroka (Eastern Highland Province), Kundiawa (Simbu Province) (n = 90), and Cairns hospital (Queensland) where isolates from routine surveillance of communities on the Cape York Peninsula were collected (n = 70). Inset schematic of the dual-panel barcoding protocol from cultured samples, where a standard spin-column extraction is followed by sequencing two rapid barcode libraries (12 isolates) on the MinION (24 samples per flow cell, interspersed with a nuclease wash) and `Sketchy` genotyping for rapid outbreak surveillance.

`Sketchy` predictions on 1000 reads of the outbreak data from FNQ and PNG were compared with Illumina reference genotypes and standard performance metrics were computed across the dataset (Figure 3, Table 2). Generally, lineage-wide distributed features (MLST, PVL, penicillin resistance, some antibiotic susceptibilities) achieved high accuracy and precision (Table 2). Importantly, most sequence types were correctly identified (144/160) providing an accurate survey of lineage diversity in FNQ and PNG. Analysis of sequence typing errors revealed consistent false calls between ST81 reference and ST1 prediction pairs (n = 3), ST243 and ST30/ST3452 (n = 5), ST5 and ST225/ST228 (n = 2) and one ST762 genome predicted as ST1 (n = 1) (Table S1). In all cases except one, the reference sequence type was contained in the reference sketch and predicted sequence types belonged to the same clonal complex (CC) with a difference of one or two alleles (Table S1). Finally, of the novel sequence types detected in this dataset (n = 5), three single allele variants of ST93 were predicted ST93, one single allele variant of ST88 was predicted ST88 and one unknown ST variant with four alleles difference was predicted ST1 (Table 2).

**Table 2.**
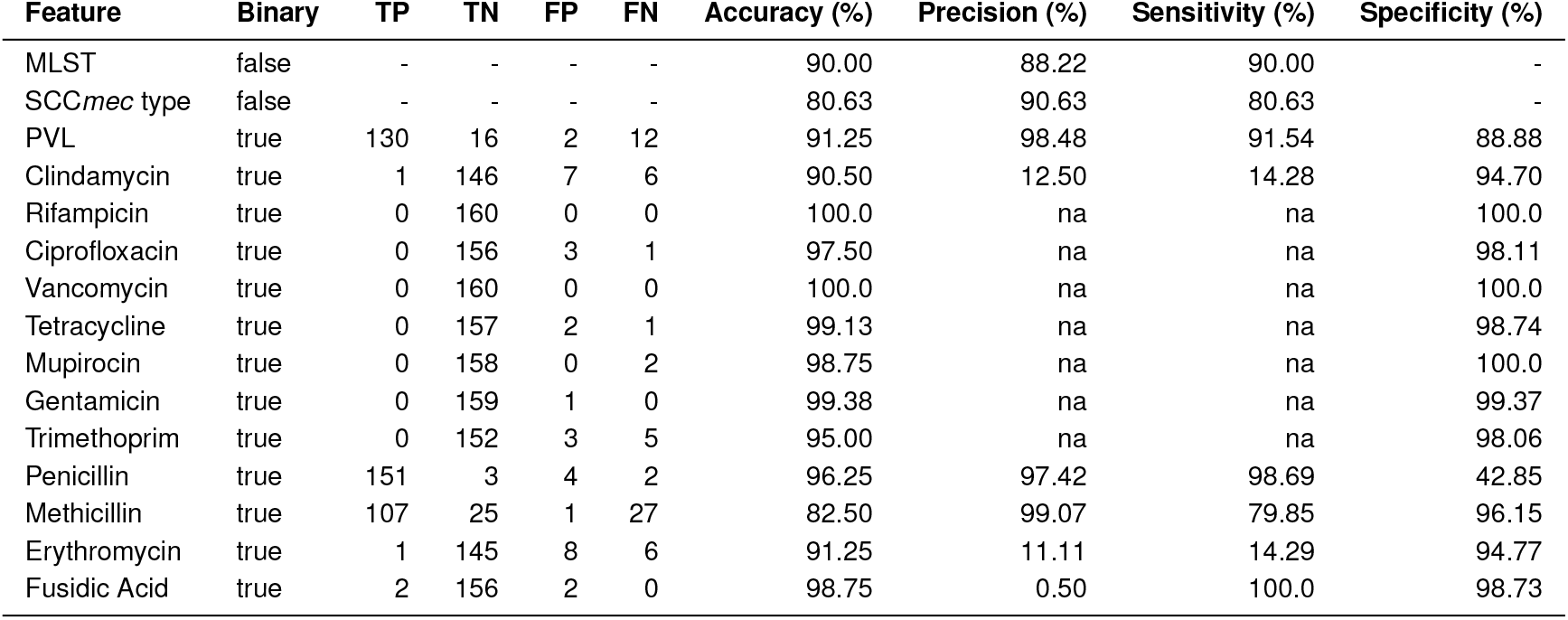
*S. aureus* outbreak isolates on MinION (4 flow cells, 2 x 12-plex, 1000 reads, k = 16, s = 1000, n = 160)

We also used the streaming algorithm (sum of shared hashes) with the same configurations for comparison (Table S2). Overall, there were slight regressions in all predictions, which were expected due to computing shared hashes per read, losing some of the information contained in the completed read sets (1000 reads). When we used the high-resolution species sketch (*s* = 10000) in non-streaming mode for these outbreak data, improvements in lineage predictions were observed, including resolution of the ST243/ST30 and ST5/ST255/ST228 misclassification, and some of the associated resistance and PVL classifications (Table S3). No improvements were made in SCC*mec* related features, with regressions in methicillin resistance and SCC*mec* type classification accuracy across the dataset, indicating that these were driven by systematic error in classification of clade-specific features of the dominant outbreak lineage ST93.

We next ran a library of strains from the osteomyelitis outbreak *in situ* at the Papua New Guinea Institute of Medical Research in Goroka (Eastern Highlands). We multiplexed 12 strains onto a MinION flow cell, but ultimately obtained few reads per barcode (276 - 1896, Table 3) due to malfunctioning laboratory equipment resulting in failed barcode attachment (15072/23774 reads unclassified, Fig. 4A). Nevertheless, we were able to use the remaining reads per barcode to type with `Sketchy`, which correctly predicted lineage (ST93) with the exception of two isolates (ST22 and ST121 from 276 and 533 reads respectively) and some lineage-distributed genotypes (PVL, some antibiotic resistance categories) suggesting deficits in SCC*mec* related predictions for ST93-MRSA-IV (Table 4). Incorrect predictions were however mitigated when the high-resolution (*s* = 10000) *S. aureus* reference sketch was used, which correctly predicted all sequence types from as few as 276 reads, but failed to predict several important secondary resistances (clindamycin, tetracycline, erythromycin) in one isolate (PNG-69) and failed to predict methicillin resistant genotypes in two others (Table S4).

**Fig. 4.**
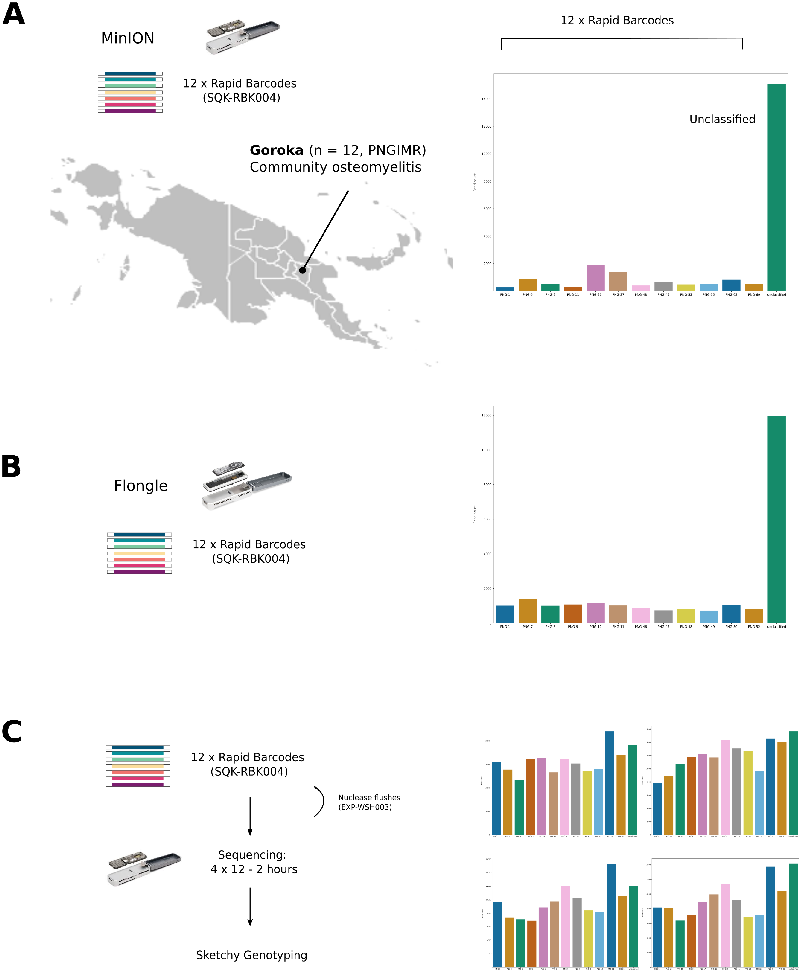
Outbreak surveillance experiments of community-associated *Staphylococcus aureus* cases from Papua New Guinea **(A)** *in situ* at the Papua New Guinea Institute for Medical Research **(B)** minimal multiplexing experiment on Flongle and **(C)** 48 strains on four successive panels on a single MinION flow cell. Left panel shows a representation of the experiment, middle panel shows the barcode distribution of each sequenced barcoded run (seagreen is unclassified). A large numbers of unclassified barcodes in **(A)** was likely due to a malfunctioning instrument (heatblock) during library preparation in Goroka, although we could not assuredly rule out other causes.

**Table 3.**
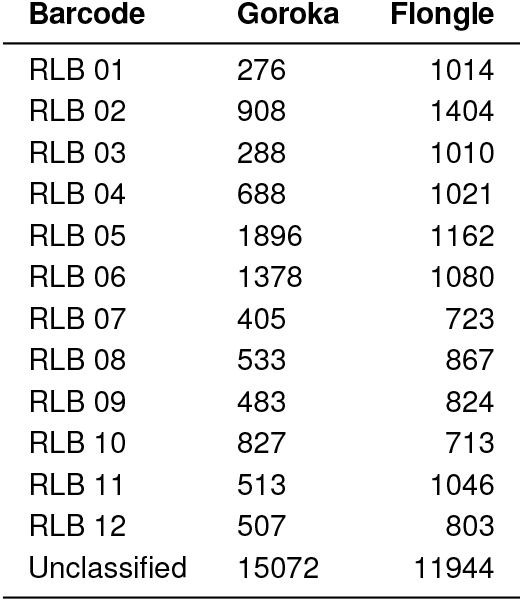
Barcode read counts

| Barcode | Goroka | Flongle |
| --- | --- | --- |
| RLB 01 | 276 | 1014 |
| RLB 02 | 908 | 1404 |
| RLB 03 | 288 | 1010 |
| RLB 04 | 688 | 1021 |
| RLB 05 | 1896 | 1162 |
| RLB 06 | 1378 | 1080 |
| RLB 07 | 405 | 723 |
| RLB 08 | 533 | 867 |
| RLB 09 | 483 | 824 |
| RLB 10 | 827 | 713 |
| RLB 11 | 513 | 1046 |
| RLB 12 | 507 | 803 |
| Unclassified | 15072 | 11944 |

**Table 4.** *S. aureus* sequencing *in situ* (Goroka) on MinION (1 flow cell, 12-plex, 1000 reads, k = 16, s = 1000, n = 12)

| Feature | Binary | TP | TN | FP | FN | Accuracy (%) | Precision (%) | Sensitivity (%) | Specificity (%) |
| --- | --- | --- | --- | --- | --- | --- | --- | --- | --- |
| MLST | false | - | - | - | - | 83.33 | 100.0 | 83.33 | - |
| SCCmec type | false | - | - | - | - | 58.33 | 100.0 | 58.33 | - |
| PVL | true | 10 | 0 | 0 | 2 | 83.33 | 100.0 | 83.33 | na |
| Clindamycin | true | 0 | 9 | 2 | 1 | 75.00 | na | na | 81.81 |
| Rifampicin | true | 0 | 12 | 0 | 0 | 100.0 | na | na | 100.0 |
| Ciprofloxacin | true | 0 | 11 | 1 | 0 | 91.66 | na | na | 91.66 |
| Vancomycin | true | 0 | 12 | 0 | 0 | 100.0 | na | na | 100.0 |
| Tetracycline | true | 0 | 11 | 0 | 1 | 91.66 | na | na | 100.0 |
| Mupirocin | true | 0 | 12 | 0 | 0 | 100.0 | na | na | 100.0 |
| Gentamicin | true | 0 | 12 | 0 | 0 | 100.0 | na | na | 100.0 |
| Trimethoprim | true | 0 | 12 | 0 | 0 | 100.0 | na | na | 100.0 |
| Penicillin | true | 12 | 0 | 0 | 0 | 100.0 | 100.0 | 100.0 | na |
| Methicillin | true | 9 | 0 | 0 | 3 | 75.00 | 100.0 | 75.00 | na |
| Erythromycin | true | 0 | 9 | 2 | 1 | 75.00 | na | na | 81.81 |
| Fusidic Acid | true | 0 | 12 | 0 | 0 | 100.0 | na | na | 100.0 |

In the next experiment we used a panel of 12 outbreak isolates to multiplex on a Flongle adapter flow cell (Fig. 4B). A large proportion of reads (11944/23611) was unclassified, which was likely due to including reads with average qualities over Q5 to account for the low throughput of the Flongle flow cell (Table 3). Similar prediction patterns as in the experiment in Goroka were observed, with few reads available for prediction across barcodes (713-1162, Table 3). Despite the low read counts, classification using the default resolution reference sketch (*k* =16, *s* = 1000, 1000 reads) successfully typed most isolates as the outbreak sequence type ST93-MRSA-IV, albeit with the previously observed limitations in SCC*mec*-related features (Table 5), including improvements across Flongle predictions with the higher resolution *S. aureus* reference sketch (Table S5). Finally, we tested a faster, successive library sequencing protocol for MinION flow cells, using 48 strains in 4 barcoded libraries, which were sequenced for 2 hours followed by a nuclease wash in between libraries (Methods). We had aimed to sequence another 4 libraries on the same flow cell (n = 96) as the 96 barcode sequencing kits had not been released yet. However, during reloading too many bubbles were introduced to the flow cell channel and the experiment terminated at 48 strains with a remaining 900-1000 active pores after a final diagnostic check (data not shown). Nevertheless, predictions of the 48 strain protocol using the default *S. aureus* sketch show that this approach is viable, with 2/48 lineage misclassifications (PNG-4, PNG-68) which were novel allele variants of ST81 and ST93 (and misclassified as ST93) (Table S1, Table 6, Table S6).

**Table 5.** *S. aureus* outbreak isolates on Flongle (1 flow cell, 12-plex, 1000 reads, k = 16, s = 1000, n = 12)

| Feature | Binary | TP | TN | FP | FN | Accuracy (%) | Precision (%) | Sensitivity (%) | Specificity (%) |
| --- | --- | --- | --- | --- | --- | --- | --- | --- | --- |
| MLST | false | - | - | - | - | 83.33 | 100.0 | 83.33 | - |
| SCCmec type | false | - | - | - | - | 75.00 | 100.0 | 75.00 | - |
| PVL | true | 10 | 0 | 0 | 2 | 83.33 | 100.0 | 83.33 | na |
| Clindamycin | true | 0 | 11 | 1 | 0 | 91.66 | na | na | 91.66 |
| Rifampicin | true | 0 | 12 | 0 | 0 | 100.0 | na | na | 100.0 |
| Ciprofloxacin | true | 0 | 12 | 0 | 0 | 100.0 | na | na | 100.0 |
| Vancomycin | true | 0 | 12 | 0 | 0 | 100.0 | na | na | 100.0 |
| Tetracycline | true | 0 | 12 | 0 | 0 | 100.0 | na | na | 100.0 |
| Mupirocin | true | 0 | 12 | 0 | 0 | 100.0 | na | na | 100.0 |
| Gentamicin | true | 0 | 12 | 0 | 0 | 100.0 | na | na | 100.0 |
| Trimethoprim | true | 0 | 12 | 0 | 0 | 100.0 | na | na | 100.0 |
| Penicillin | true | 12 | 0 | 0 | 0 | 100.0 | 100.0 | 100.0 | na |
| Methicillin | true | 9 | 0 | 0 | 3 | 75.00 | 100.0 | 75.00 | na |
| Erythromycin | true | 0 | 9 | 2 | 1 | 91.66 | na | na | 91.66 |
| Fusidic Acid | true | 0 | 12 | 0 | 0 | 100.0 | na | na | 100.0 |

**Table 6.** Successive library experiment on MinION (1 flow cell, 4 x 12-plex, 1000 reads, k = 16, s = 1000, n = 48)

| Feature | Binary | TP | TN | FP | FN | Accuracy (%) | Precision (%) | Sensitivity (%) | Specificity (%) |
| --- | --- | --- | --- | --- | --- | --- | --- | --- | --- |
| MLST | false | - | - | - | - | 95.83 | 91.84 | 95.83 | - |
| SCCmec type | false | - | - | - | - | 87.50 | 94.24 | 87.50 | - |
| PVL | true | 46 | 0 | 1 | 1 | 95.83 | 97.87 | 97.87 | na |
| Clindamycin | true | 1 | 47 | 0 | 0 | 100.0 | 100.0 | 100.0 | 100.0 |
| Rifampicin | true | 0 | 48 | 0 | 0 | 100.0 | na | na | 100.0 |
| Ciprofloxacin | true | 0 | 46 | 1 | 1 | 95.83 | na | na | 97.87 |
| Vancomycin | true | 0 | 48 | 0 | 0 | 100.0 | na | na | 100.0 |
| Tetracycline | true | 0 | 47 | 0 | 1 | 97.91 | na | na | 100.0 |
| Mupirocin | true | 0 | 48 | 0 | 0 | 100.0 | na | na | 100.0 |
| Gentamicin | true | 0 | 48 | 0 | 0 | 100.0 | na | na | 100.0 |
| Trimethoprim | true | 0 | 48 | 0 | 0 | 100.0 | na | na | 100.0 |
| Penicillin | true | 48 | 0 | 0 | 0 | 100.0 | 100.0 | 100.0 | na |
| Methicillin | true | 41 | 1 | 1 | 5 | 87.50 | 97.61 | 89.13 | 0.50 |
| Erythromycin | true | 1 | 47 | 0 | 0 | 100.0 | 100.0 | 100.0 | 100.0 |
| Fusidic Acid | true | 0 | 48 | 0 | 0 | 100.0 | na | na | 100.0 |

### Sublineage genotyping comparison with RASE

Finally, we compared `Sketchy` at sublineage resolution to `RASE` predictions for the outbreak sequence type (ST93). We built reference databases based on lineage genomes (n = 360, *k* =16) including a rooted maximum-likelihood phylogeny from previous single nucleotide polymorphism calls (26) (Fig. 5A). Because of the small size of this reference databases, we constructed additional sketches with higher resolution (up to *s* = 1000000) to compare for sublineage genotyping with `Sketchy` (Methods). `RASE` predictions were largely congruent with reference genotypes, with most categories exceeding 90% accuracy and precision, and only sporadic false positive and false negative predictions for clindamycin, mupirocin, methicillin and erythromycin (Table 7). There appeared to be a systematic error in tetracycline predictions, where 28/118 isolates were predicted resistant (R), but were in fact susceptible (S). Only a single isolate assembly in the reference database was typed as resistant (R). We ruled out contaminated genomes in the reference sketch as a source for these aberrant predictions, due to using conservative filters including contamination and strain heterogeneity (Methods). In addition, we ruled out errors introduced by ancestral state reconstruction, which was disabled for this analysis in `RASE`. Ultimately, most false tetracycline resistance predictions were flagged with low confidence from the preference score used in `RASE`, but did not resolve when using all reads for inference (Table S5).

**Fig. 5.**
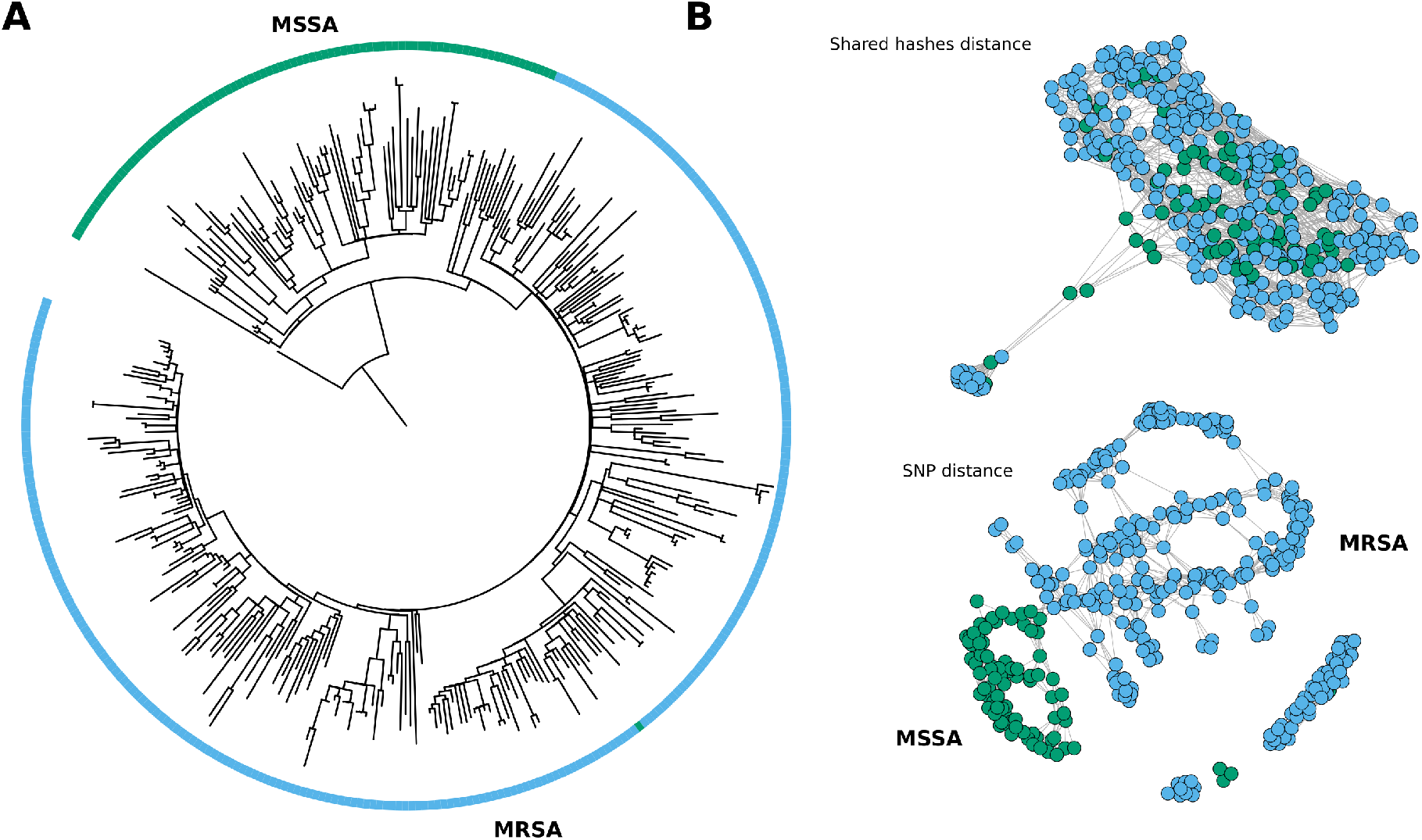
Phylogenetic tree for sublineage genotyping comparison with `RASE` and visualization of population structure with `NetView` **(A)** Maximum-likelihood phylogeny of the ST93 reference genomes (n = 360) used in `RASE` showing the ancestral MSSA clade (green) and the divergent MRSA clade (blue). **B** Mutual k-nearest-neighbor graphs (`NetView`) for visualization of MSSA/MRSA population structure using the shared hashes distance computed with `Sketchy` (*k* =16, *s* = 1000) failing to distinguish between the two genotype clades and SNP distances underlying the ML phylogeny, successfully resolving the MSSA/MRSA clades. Differences between the two core methods (shared hashes and SNPs) represent the limitations of `Sketchy` to predict methicillin resistance genotypes at sublineage resolution; homogeneous network topologies for shared hashes distances are obtained for *s* = 1000 – 1000000 (data not shown).

**Table 7.** `RASE` classification of ST93 outbreak isolates (n = 120, lineage database)

| Feature | Binary | TP | TN | FP | FN | Accuracy (%) | Precision (%) | Sensitivity (%) | Specificity (%) |
| --- | --- | --- | --- | --- | --- | --- | --- | --- | --- |
| Clindamycin | true | 0 | 113 | 4 | 3 | 94.16 | na | na | 96.58 |
| Rifampicin | true | 0 | 120 | 0 | 0 | 100.0 | na | na | 100.0 |
| Ciprofloxacin | true | 0 | 120 | 0 | 0 | 100.0 | na | na | 100.0 |
| Vancomycin | true | 0 | 120 | 0 | 0 | 100.0 | na | na | 100.0 |
| Tetracycline | true | 1 | 90 | 29 | 0 | 75.83 | 3.33 | 100.0 | 75.63 |
| Mupirocin | true | 0 | 115 | 5 | 0 | 95.83 | na | na | 95.83 |
| Gentamicin | true | 0 | 120 | 0 | 0 | 100.0 | na | na | 100.0 |
| Trimethoprim | true | 0 | 120 | 0 | 0 | 100.0 | na | na | 100.0 |
| Penicillin | true | 120 | 0 | 0 | 0 | 100.0 | 100.0 | 100.0 | na |
| Methicillin | true | 110 | 0 | 3 | 7 | 91.66 | 97.34 | 94.02 | na |
| Erythromycin | true | 0 | 113 | 4 | 3 | 94.16 | na | na | 96.58 |
| Fusidic Acid | true | 0 | 120 | 0 | 0 | 100.0 | na | na | 100.0 |

`Sketchy` performed slightly worse than `RASE` using a low resolution sketch (*s* = 1000) (Table 8) with sporadic false positives and false negatives in clindamycin, ciprofloxacin, tetracycline, mupirocin and erythromycin predictions. However, these were largely eliminated using the high-resolution sketch (*s* = 1000000) raising accuracy and precision for most antibiotic resistance predictions to *>* 96% (Table 9). `RASE` timestamps indicate that predictions of ST93 genotypes around the selected read threshold (1000 reads) were able to be conducted in approximately 1 - 11 minutes per barcode (data not shown). According to our expectations, systematic errors were found in the methicillin predictions of `Sketchy`, with an excess of isolates that were typed susceptible (MSSA) rather than resistant (MRSA). `Sketchy` was therefore not capable of sufficiently resolving clade-specific traits for sublineage genotyping. We illustrated the difference in resolution of the underlying core method (MinHash vs. SNPs) and its ability to resolve clade-specific traits in the ST93 reference sketch using population graphs, where nodes are genomes and edges their mutual *k*-nearest-neighbors at an optimized *k* value (genomic neighborhoods) (Fig. S1D). We constructed the graph for pairwise *s* - shared hashes distance (*s* = 1000) using `Sketchy` as well as from pairwise SNP distances based on previously generated variants for the ST93 lineage (26) (from which the ML topology in the `RASE` approach was built) (Fig 5B). Shared hash distances were insufficient to resolve MSSA and MRSA communities compared to networks constructed from pairwise SNP distances. This fundamental difference in resolution of the two approaches underlines the limitations of `Sketchy`, although the ultra high-resolution sketch (*s* = 1000000) mitigated some of the non clade-specific errors (e.g. clindamycin and erythromycin resistance) observed using lower-resolution sketches (Table 9).

**Table 8.**
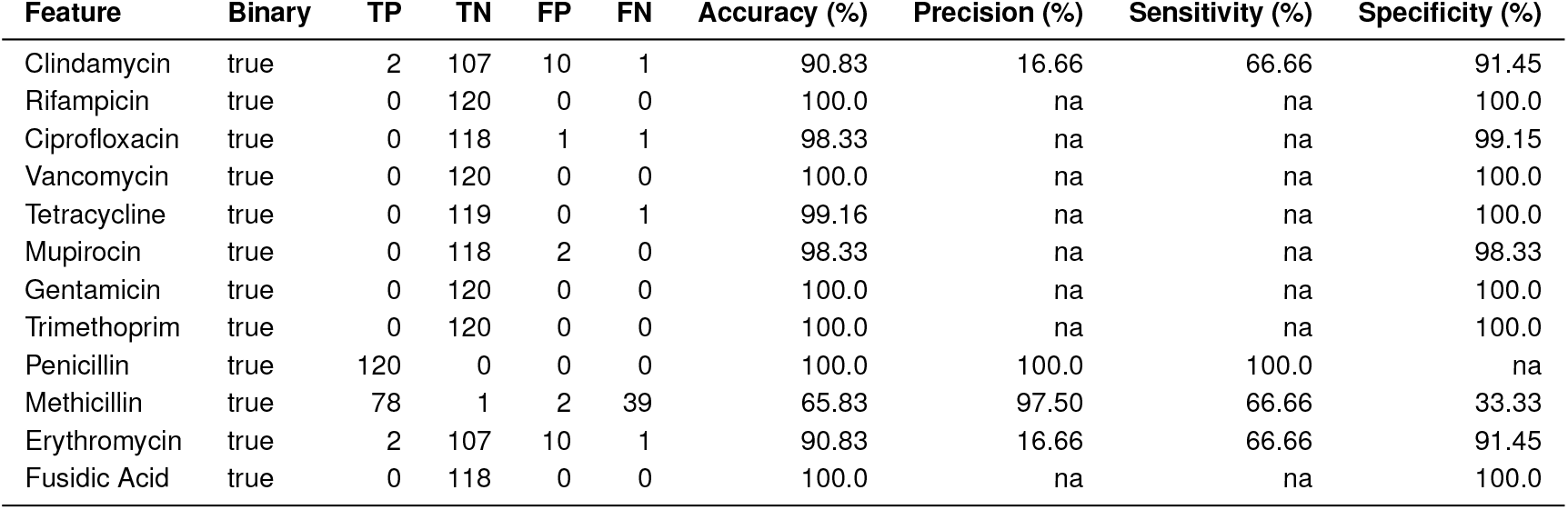
`Sketchy` classification of ST93 outbreak isolates (n = 120, lineage sketch, s = 1000)

**Table 9.** `Sketchy` classification of ST93 outbreak isolates (n = 120, lineage sketch, s = 1000000)

| Feature | Binary | TP | TN | FP | FN | Accuracy (%) | Precision (%) | Sensitivity (%) | Specificity (%) |
| --- | --- | --- | --- | --- | --- | --- | --- | --- | --- |
| Clindamycin | true | 0 | 116 | 1 | 3 | 96.66 | na | na | 99.14 |
| Rifampicin | true | 0 | 120 | 0 | 0 | 100.0 | na | na | 100.0 |
| Ciprofloxacin | true | 0 | 119 | 0 | 1 | 99.16 | na | na | 100.0 |
| Vancomycin | true | 0 | 120 | 0 | 0 | 100.0 | na | na | 100.0 |
| Tetracycline | true | 0 | 119 | 0 | 1 | 99.16 | na | na | 100.0 |
| Mupirocin | true | 0 | 120 | 0 | 0 | 100.0 | na | na | 100.0 |
| Gentamicin | true | 0 | 120 | 0 | 0 | 100.0 | na | na | 100.0 |
| Trimethoprim | true | 0 | 120 | 0 | 0 | 100.0 | na | na | 100.0 |
| Penicillin | true | 120 | 0 | 0 | 0 | 100.0 | 100.0 | 100.0 | na |
| Methicillin | true | 87 | 1 | 2 | 30 | 73.33 | 97.75 | 74.35 | 33.33 |
| Erythromycin | true | 0 | 116 | 1 | 3 | 96.67 | na | na | 99.14 |
| Fusidic Acid | true | 0 | 118 | 0 | 0 | 100.0 | na | na | 100.0 |

## Discussion

In this study, we explored the use of heuristic genomic neighbor typing (16) for lineage and genotype inference in bacterial outbreak scenarios. We reasoned that a ‘hypothesis-agnostic’ reference database would be preferred over a smaller ‘hypothesis-driven’ reference database, because the latter cannot capture the known diversity of a species, and may not be useful in situations where prior sequence data on lineage and genotype diversity does not exist. We further reasoned that it would be possible to conduct multiplex sequencing and use genomic neighbor typing to rapidly scan an isolate collection from limited sequence data. For these applications we developed `Sketchy`, a genomic neighbor typing implementation using shared min-wise hashes against species-wide database sketches with associated lineage and genotype data, which we derived from public sources (25).

We first used a cross-validation procedure to assess performance of `Sketchy` at the selected 1000 reads threshold in recovering lineages (MLST) as a proxy for further sublineage genotyping (Fig. 2, Table 1) using default resolution (*s* = 1000) and higher resolution (*s* = 10000) reference sketches. Results indicate two major trends: first, genomic neighbor typing of lineages was sufficiently accurate in some species (*S. aureus*, *K. pneumoniae*, *P. aeruginosa*) but failed to recover lineages in others (*N. gonorrhoeae*, *N. meningitidis*, *S. pneumoniae*). We were unable to control for sequence type diversity of the reference sketches, and it may be possible that cross-validation sampling of reference sketches with many singular sequence types or little sequence type diversity biased the results in favour of sketches with fewer diversity. However, we note that the species which did not perform well on this task are those with high levels of homologous recombination. This was discussed by Břinda *et al*. (16) who suggested that genomic neighbor typing may be limited by homologous recombination due to scattering of the phylogenetic signal and spread of chromosomally encoded resistance genes. We note that `Sketchy` under-performed for MLST typing of two species, *N. gonorrhoeae* and *S. pneumoniae*, both of which were used in the original genomic neighbor typing approach. However, direct comparisons are difficult, as the underlying reference data was vastly smaller, and the focus was on sublineage antimicrobial resistance phenotyping using MIC values from the reference collection.

Second, we observed that performance notably increased using a larger sketch size, suggesting that the default sketch size may be insufficient to capture the full diversity of hashes shared between analyte and the reference databases. This suggests that the default sketch size (*s* = 1000) may not be a good default if accuracy is preferred. However, because memory consumption increased approximately linearly with the number of included genomes and sketch size, higher resolution sketches may not be suitable for smaller computing platforms, especially with large reference databases (e.g. for *S. aureus* and *S. pneumoniae*). Nevertheless, memory consumption did not exceed 6 GB, making large, species-wide reference sketches at higher resolution usable on laptops and other standard computing hardware. In addition, we note that memory consumption of `Sketchy` for the smallest reference sketch (*P. aeruginosa*, 4832 genomes, 56 MB) is significantly smaller than the `ProPhyle` (28) indices created for *S. pneumoniae* (616 genomes, 321 MB) and *N. gonorrhoeae* (1102 genomes, 242 MB), and for higher resolution sketches approximately twice as much (554 MB) (29). Overall, sketch sizes are extremely small, particularly when compressed for transfer or storage (Table 1). `Sketchy` is therefore capable of creating highly efficient species-wide databases, which can capture the known diversity of a species, while maintaining resource efficiency, albeit with some limitations in performance for smaller sketches which may be necessary for portable sequencing setups in remote locations.

We then assessed `Sketchy`’s performance on an outbreak dataset of *S. aureus* community infections in Papua New Guinea and Far North Queensland (4, 17, 18), for which we had previously generated matching Illumina reference data (n = 160). In this context, the outbreaks constituted an independent validation dataset, as no *S. aureus* genomes from these regions had ever been sequenced before (including the very first *S. aureus* genomes from Papua New Guinea). While the majority of isolates in this study belonged to the outbreak sequence type (ST93, n = 120) several other sequence types were identified in the dataset (n = 40). By including all known strains at the time in the reference database, their lineages were included during database construction by default and successfully typed in most cases (Table 4). Remaining misclassifications were mitigated in higher resolution sketches (Table S3) with the exception of clade-specific *SCCmec* related genotypes (see below). In addition, we sequenced one full panel of outbreak isolates at the Papua New Guinea Institute of Medical Research in Goroka, Eastern Highlands Province. No sequencing infrastructure is accessible, so that a portable setup with the MinION was the only option to survey the outbreak on site. As an illustration of the challenges of sequencing in remote places, a heatblock malfunctioned during library preparation, which was likely the reason for sub-optimal barcode attachment resulting in extremely low throughput for a MinION flow cell (Table 3). Nevertheless, we were able to obtain 83% (default resolution) and 100% accuracy (higher resolution, Table S4) in typing lineages, providing a useful picture of the outbreak sequence type, antibiotic resistance genotypes (with exception of SCC*mec* related features) and presence of the PVL toxin. Similar results were obtained on a multiplex run on cheap Flongle adapters (Table 5, Table S5).

We employed an efficient multiplex sequencing protocol on the MinION for surveying the two outbreaks, sequencing 2 x 12 barcodes on the same flow cell, driving down the cost of each isolate with full assembly, genotyping and phylodynamic analysis to around $40-50 per isolate, as previously described (4). In this analysis, we used a subset of the total reads per barcode (1000) for genomic neighbor typing evaluation which corresponds to approximately 2-3x coverage of the *S. aureus* genome, having shown previously that assembly based genotyping is possible at approximately 5x coverage per genome (4). In addition, we expanded on the dual-library sequencing protocol and attempted to sequence 48 strains on a MinION flow cell in 2 hour intervals, with sufficient data obtained for genomic neighbor typing further reducing cost to approximately $30 (Australian) per barcode (Table 6, Table S6). Given the efficiency of our approach, it should be possible to use 96-barcode kits to sequence as many isolates on a single MinION flow cell and obtain accurate genotypes. Taking into consideration the limitations in species applications and resource management for higher resolution sketches, genomic neighbor typing with `Sketchy` is therefore suitable to survey bacterial outbreaks rapidly, at low-cost, and with sufficiently accurate results to infer important epidemiological characteristics. In this case, the predominant outbreak sequence type was the Australian ST93 lineage, which had emerged in the Northern Territory and spread to the East Coast of Australia (26). Even without confirmation from phylogenetic analysis (27), the predominance of the ST93 sequence type in Far North Queensland and Papua New Guinea outbreaks strongly suggests transmission from Australia.

Finally, we observed systematic misclassifications of clade-specific SCC*mec* related features (methicillin resistance, SCC*mec* subtype) that could not be resolved with higher resolution sketches (Tables 2–6, Tables 8–9). We hypothesized that this could be due to the approximate MinHash approach, which does not have the same resolution on sublineage geno- or phenotypes as the phylogenetically guided classification approach using `ProPhyle` in `RASE`. We demonstrated this limitation on a lineage-specific (ST93) reference sketch in comparison with the same reference database implemented in `RASE` (Tables 7–9), for which we used a phylogenetic tree of the lineage that distinguished between MSSA and MRSA clades (Fig. 5A). While most misclassifications with `Sketchy`could be resolved with increasing sketch size (Table 9) and indeed outperformed `RASE`, classifications of SCC*mec* features continued to fail even at very high sketch sizes (*s* = 1000000). While `RASE` performed better on sublineage genotyping, we noted a systematic error in tetracycline predictions, which was unexpected since only a single isolate in the reference dataset was resistant; we were unable to explain these errors but note that the preference score employed by `RASE` marked uncertainty in the majority of tetracycline predictions, even when run on all reads for each isolate, ultimately not resolving the tetracycline prediction errors (Table S5).

Overall, phylogenetically informed genomic neighbor typing has a definitive advantage over `Sketchy` for inference of clade-specific traits, which his particularly relevant for clinical diagnostics (e.g. antimicrobial susceptibility predictions). However, we were unable to construct `RASE` databases for the species-wide reference collections, as the required phylogenetic trees are infeasible, or at least highly impractical, to infer from tens of thousands of whole genome sequences. At the species level, the ease with which reference sketches can be constructed for *Sketchy* and their minimal resource requirements given the number of genomes included, puts `Sketchy` at an advantage for outbreak surveillance applications. Because we derive genotypes from other genotype classifications (based on assemblies or reads) it should be noted that classification with `Sketchy` can only achieve classification performance of the underlying genotyping methods (e.g. `Mykrobe`). However, genomic neighbor genotyping with `Sketchy` could also enable automating the construction of reference databases, so that public archives can be surveyed periodically and new genome integrated continuously. At this stage, due to limitations in sublineage genotype predictions for antibiotic susceptibility predictions, we do not recommend using `Sketchy` for clinical applications, but rather as a tool to rapidly survey bacterial outbreaks or isolate collections at scale. `Sketchy` may also be useful in scenarios where genotype inference of limited sequence data is required.

In comparison to other genotype classification tools, `Sketchy` is situated between species-level taxonomic classifiers and phylogenetically informed genomic neighbor typing. Predictions are useful for traits distributed at the lineage level, for example penicillin resistance or PVL toxins in *S. aureus*, with limitations in application to some species with high rates of homologous recombination, such as *Neisseria gonorrhoeae* or *Neisseria meningitidis*. Future work on genomic neighbor typing may consider scaling up multiplexing (e.g. 96-barcode panels), curation and minimisation of reference databases, implementation of alternative query methods, or combining different approaches to genomic neighbor typing to enable continuous species- to sublineage-level predictions. Adaptive sequencing may be useful to balance throughput per barcode in order to make multiplex sequencing protocols more robust and cost-effective (30). Ultimately, we demonstrated that genomic neighbor typing with species-wide reference sketches is a viable approach for genotype surveillance of bacterial community outbreaks, particularly under challenging circumstances and in remote locations, including northern Australia and Papua New Guinea.

## Materials and Methods

### Outbreak sampling and reference sequencing

Isolates were collected from outbreaks in two remote populations in northern Australia and Papua New Guinea as described by Steinig et al. (4) and Aglua et al. (17). Isolates associated with paediatric osteomyelitis cases (mean age of 8 years) were collected from 2012 to 2017 (n = 42) from Kundiawa, Simbu Province (27), and from 2012 to 2018 (n = 35) from patients in the neighboring Eastern Highlands province town of Goroka. We supplemented the data with MSSA isolates associated with severe hospital-associated infections and blood cultures in Madang (Madang Province) (n = 8) and Goroka (n = 12). Isolates from communities in Far North Queensland, including metropolitan Cairns, the Cape York Peninsula and the Torres Strait Islands (n = 91) were a contemporary sample from 2019. Isolates were recovered on LB agar from clinical specimens using routine microbiological techniques at Queensland Health and the Papua New Guinea Institute of Medical Research (PNGIMR). Isolates were transported on swabs from monocultures to the Australian Institute of Tropical Health and Medicine (AITHM Townsville) where they were cultured in 10 ml LB broth at 37°C overnight and stored at −80°C in 20% (v/v) glycosol and LB. Samples were regrown on LB agar prior to sequencing, and a single colony was placed into inhouse lysis buffer and sequenced at the Doherty Applied Microbial Genomics laboratory (DAMG), using 100 bp paired-end libraries on Illumina HiSeq.

### MinION outbreak library preparation and sequencing

2 ml of LB broth was spun down at 5,000 x g for 10 minutes and after removing the supernatant, 50 ul of 0.5 mg / ml lysostaphin were added to the tube and vortexed. Cell lysis was conducted at 37°C for 2 hours with gentle shaking followed by a *proteinase K* digestion for 30 mins. at 56°C. DNA was extracted using a simple column protocol from the DNeasy Blood & Tissue kit (QIAGEN) following the manufacturer’s instructions. DNA was eluted in 70 ul of nuclease-free water, quantified on Qubit, and DNA was stored at 4°C until library preparation. Library preparation was done using approx. 420 ng of DNA and the rapid barcoding kit with 12 barcodes (ONT, SQK-RBK004) as per manufacturer’s instructions, with the exception of conducting bead cleanup steps. DNA was quantitated using Qubit 4.0 (Thermo Fisher Scientific), purity determined with a NanoDrop 2000 Spectrophotometer (Thermo Fisher Scientific). Basecalling was done using the PyTorch Bonito v0.3.6 R9.4.1 DNA model, run on a local NVIDIA GTX1080-Ti or a remote cluster of NVIDIA P100 GPUs. Sequence runs were conducted with 2 x 12 barcoded (SQK-RBK004) isolates per flow cell in two consecutive 18-24 hour runs. Libraries were nuclease flushed using the wash kit between consecutive runs (Oxford Nanopore Technologies, EXP-WSH-003). This is sufficiently effective to remove read carry-over, as demonstrated previously with hybrid assemblies of sequentially sequenced *Enterobacteriaceae* (31). Sequencing runs were managed on two MinIONs and monitored in MinKNOW > v20.3.1. Read sum mary reports for nanopore reads were generated with nanoq v0.8.2 (32).

### MinION and Flongle multiplexing experiments

To demonstrate that genotyping is possible on site in Papua New Guinea, we sequenced 12 *S. aureus* outbreak strains at the Papua New Guinea Institute of Medical Research (PNGIMR) in Goroka. We replicated the QIAGEN extraction and rapid library sequencing protocol described above, unknowingly using a malfunctioning heatblock in the library preparation step (SQK-RBK004), which resulted in suboptimal barcode attachments. We also prepared a multiplex run for a Flongle experiment at the Peter Doherty Institute for Infection and Immunity. *Staphylococcus aureus* glycerol stocks were inoculated in Tryptic soy broth (TSB) and grown overnight at 37°C, 180 rpm. DNA was extracted from 8 ml of overnight culture via pelleting cells at 12,000 rpm for 2 minutes. Cells were resuspended in PrepMan™ Ultra Sample Preparation Reagent (ThermoFisher Scientific) and Lysing Matrix Y beads (MP Biomedicals). Isolates were incubated at 95°C for 15 minutes and cells further lysed via a TissueLyser LT (Qiagen) at 6.5 m/s for 60 seconds similar to previously described (33). Extracts were centrifuged at 13,000 rpm for 10 minutes. Supernatant was removed and mixed with 3M sodium acetate (pH 5.5), ice-cold 100% ethanol (0.3:0.03:0.67 ratio) and DNA was precipitated for 3 hours at −20°C. DNA was pelleted at 13,000 rpm for 15 mins (4°C), washed with 70% ethanol and resuspendeded in ultrapure water. High-molecular-weight (HMW) DNA was isolated via the MagAttract HMW DNA Kit (Qiagen) as per manufacturer’s instructions. Briefly, this included a protein digest with proteinase K for 30 minutes at 56°C (900 rpm) and an RNAse A (0.4mg) treatment for 10 minutes at room temperature. HMW DNA was further purified using Agencourt Ampure XP (Beckman Coulter Australia) beads (1:1 ratio). Libraries were prepared using the ONT Rapid Barcoding (SQK-RBK004) kit with an input of 200ng of HMW DNA for each isolate. The library was sequenced on an ONT Flongle FLO-FLG001 flow cell for 24 hours. All runs in this sections were called with `Guppy` v4.6 R9.4.1 DNA high accuracy models (HAC). Finally, we repeated library construction as described for the outbreak sequencing above to test a faster sequencing protocol, in which four libraries were sequenced on the same MinION flow-cell with intermediate nuclease flushes and a runtime of 2 hours per library.

### Reference databases construction and genotyping

For reference sketch construction, we used a collection of assemblies containing bacterial genomes from the entire European Nucleotide Archive (ENA) in 2018 (n = 660,333) (25). Metadata from precomputed assembly genotypes was used to subset assemblies with complete lineage designation for inclusion (MLST). `CheckM` metrics were used to filter assemblies by completeness (< 99%), contamination (> 0.1%) and evidence for strain heterogeneity (> 0.1%) retaining a total of 543,695 assemblies across 71 species with at least 100 genomes. For reference sketch construction in the simulations, we included five common species of interest with at least 1000 genomes: *Streptococcus pneumoniae* (n = 47,616), *Staphylococcus aureus* (n = 42,461), *Neisseria meningitidis* (n = 16,198), *Klebsiella pneumoniae* (n = 10,072) and *Neisseria gonorrhoeae* (n = 8,413). We had previously downloaded a collection of *S. aureus* sequence runs from the NCBI Short Read Archive and ENA (n = 38,985) providing matching raw sequence read data for a subset of the assemblies in the ENA collection. Antimicrobial resistance phenotypes for 12 antibiotics (ciprofloxacin, clindamycin, erythromycin, fusidic acid, gentamicin, methicillin, mupirocin, penicillin, rifampicin, tetracycline, trimethoprim and vancomycin) were inferred from these reads with `Mykrobe v0.6.1` and the default *S. aureus* typing panel (34). In addition, we used `SCCion v0.2.1` (https://github.com/esteinig/sccion) to type *SCCmec* subtypes using `Mash` matches against the SCC*mec*Finder database (35).

### Sketchy implementation and streaming algorithm

`Sketchy` implements k-mer extraction and hashing based on the `needletail` (https://github.com/onecodex/needletail) and `finch` (https://github.com/onecodex/finch-rs) libraries, which allowed us to replicate `Mash` sketching and shared hashes computation in Rust. `Mash` (23) pioneered an unbiased approximation of the Jaccard index between two k-mer sets *A* and *B*:

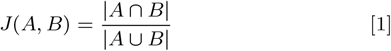

Mash (and Finch) merge-sort two bottom sketches *S*(*A*) and *S*(*B*) to estimate the Jaccard index, where the merge is terminated after *s* unique hashes, and the estimate of the Jaccard index is computed for *x* shared hashes found after processing *s*′ hashes:

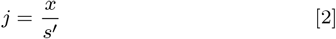

`Sketchy` implements two simple reference sketch matching functions based on the parameters of the reference sketch (k-mer size, sketch size and hash seed) that compute the min-wise shared hashes (*x*) with each genome in the reference sketch. In the first instance, we use `Finch` to compute the number of shared hashes (*x*) for all reads until the specified read limit (*i*) (`--limit` parameter). In addition, we provide a streaming implementation (the sum of shared hashes) in which the shared hashes (*x*) are computed for each read (*j*) and added to the sum of shared hashes (*h*) until the read limit (*i*) is reached:

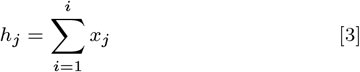

Implementation of genomic neighbor typing is achieved by ranking the shared hashes (or the sum of shared hashes after each read) and selecting the associated genotype of the highest ranking genome in the reference database as inferred genotype. When predicting genomic neighbors from closely related genomes of the same lineage (e.g. in an outbreak scenario) a consensus call for each genotype features across the highest ranking genomes can be made using the `--consensus` flag and `--top` parameter in the `Sketchy` command line client.

### Sketchy command line client

`Sketchy v0.6.0` is written in Rust and implements a command line client with several functions. First, a multi-genome reference sketch can be constructed from sequence files at a given sketch (*s*) and k-mer size (*k*):

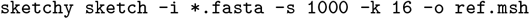

Information about the sketch (k-mer size, sketch-size, hash seed, number of genomes, identity and order of genomes) can be produced from the sketch file:

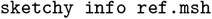

Associated genotype or phenotype files can then be constructed and checked against the reference sketch to ensure both contain the same genomes in the same order:

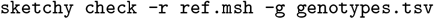

For any given (multiple) sketch file the shared hashes with each genome in the reference sketch can then be computed, if parameters between the reference and query sketches are consistent:

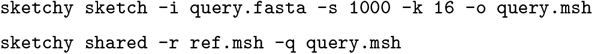

Finally, genomic neighbor typing predictions based on the reference sketch and a sequence file can be computed for a given number of reads (`--limit`), which will output a given number of the highest ranking matches (`--top`) in the reference sketch and their associated genotypes or phenotypes for inference. Streaming and consensus modes can be activated with their respective flags (`--stream` and `--consensus`):

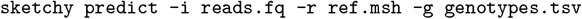

### Lineage calling simulations and comparisons

Databases varied in the representation of the total diversity within each species, due to variations in the number of genomes available and diversity of sequence types contained in each database. We conducted a cross-validation analysis by randomly sampling 10 genomes from each database across multiple replicate samples (n = 20). For each replicate, we constructed the reference sketch without the sampled genomes to evaluate the impact of predicting sequence types not contained in the database. We used `badread v0.2.0` to simulate decent quality, low-coverage (5x) nanopore reads (similar to using R9.4 flow cells and RAD004 libraries) with parameters:

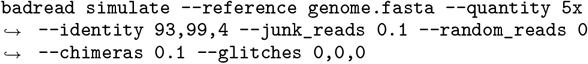

We then selected a series of read cut-offs for predictions (10, 50, 100, 200, 300, 500 and 1000 reads). Ultimately, we selected to report results at the 1000 read threshold for several reasons: first, the threshold marks around 1-3x coverage of the *S. aureus* genome (depending on read length), after which it becomes feasible to do assembly based genotyping with high recall from nanopore data alone, as demonstrated previously for these outbreak data (20); second, our primary aim was to infer genotypes from as few reads as possible and initial simulations indicated stabilisation of predictions below 1000 reads (Fig. S1); third, reporting by time (as in `RASE`) is highly volatile due to differences in throughput between libraries (e.g. multiplex vs. single isolates), sequencing devices (e.g. MinION vs. PromethION) as well as pore availability and occupancy per flow cell. Our target for these simulations was lineage calling, as the prediction of intra-lineage genotypes (including antibiotic resistance) depends on first matching into the correct genomic neighborhood of the species (i.e. finding the correct sequence type). MLST (lineage) predictions were made from the match with the highest shared hashes in the replicate database (`--top 1`). Replicate samples were run against the hold-out sketches (DB-) and against the full sketch (DB+) computing the average sequence types correctly predicted over all samples (including standard deviation, Table 1).

For comparison at the 1000 read threshold we used `Krocus v1.0.1` (*k* = 16), which attempts to find k-mers matching to species-specific MLST alleles and is conceptually similar to `Sketchy` in that it implements a ‘hypothesis-agnostic’ approach to genotyping lineages (based on available MLST alleles from PubMLST) (36). We also compared results with assemblies of the simulated genomes using `Flye v2.9` (37) followed by MLST typing with `mlst v2.19.0` (https://github.com/tseemann/mlst). At this stage, we did not compare `Sketchy` to `RASE`, because `RASE` requires phylogenetic guide trees for `ProPhyle` (28), which are not feasible or practical to infer for species-wide whole genome collections, such as the ones constructed here. Direct inference of MLST from assemblies and k-mer allele typing were therefore conceptually more suitable for comparison with `Sketchy`. Mean maximum memory consumption and time for prediction were measured on a single representative isolate picked at random for 10 iterations (including standard deviation, Table 1).

### Genotyping of community-associated outbreaks

For validation of predictions in an outbreak surveillance scenario we used a set of 160 nanopore-sequenced isolates from FNQ (n = 70) and PNG (n = 90) sequenced using the dual-library protocol and for which we had matching Illumina data. Using Illumina genotypes as reference, for each binary genotype feature (e.g. R or S, PNL+ or PVL-) we computed accuracy, precision, sensitivity, and specificity using `sklearn` functions, with weighed scores for multi-label features (SCC*mec*-type, MLST). While the dataset constituted a real test dataset with previously unknown strains from a country for which genome sequences did not exist for *S. aureus*, it should be noted that there was substantial bias in composition towards the ST93-MRSA-IV outbreak lineage (n = 120/160). Sketchy was run using consensus genotypes over the 5 highest ranking prediction of the default reference sketch for *S. aureus* (*k* = 16, *s* = 1000, 1000 reads classification limit) which marginally improved within outbreak genotyping of ST93 isolates. Output predictions were evaluated against the Illumina reference genotypes for each feature (Tables 2 – 5). For comparison of streaming analysis (sum of shared hashes) we used the outbreak dataset and the highest ranking prediction (1000 reads classification limit) (Table S3). For demonstration of applying `Sketchy` in challenging sequencing scenarios related to this outbreak, we conducted three experiments: a multiplex flow cell of 12 outbreak isolates sequenced in Goroka (during which a heatblock failed resulting in suboptimal barcode attachment), a library on an early Flongle adapter flow cell with highly reduced throughput, and sequencing 4 panels of 12 barcoded isolates in succession (2 hours each, with nuclease washes between runs, see above). We applied the same consensus genotype prediction and metrics for these three experiments as in the dual barcoding library (Fig. 4, Tables 3–5).

### Genomic neighbor typing of sublineage genotypes

For comparison of sublineage antimicrobial resistance typing with `RASE`, we collected a reference set of ST93-MSSA and -MRSA strains based on previous work with this lineage (n = 360) (26). Genotype data consisted of the antimicrobial resistance genotypes derived from the full reference sketch for *S. aureus* used in the outbreak surveillance section. For implementation in `RASE`, we constructed a phylogenetic tree based on core SNPs from our previous phylogenomic analysis of the lineage (26). `IQTREE v2.1.2` was used to reconstruct a maximum-likelihood phylogeny using the General Time Reversible model with rate heterogeneity, Lewis ascertainment bias correction (GTR+G+ASC) and placing the root on an early diverging MSSA isolate, consistent with previous phylogenetic reconstructions (SAMEA1557252). Trees were visualized with Interactive Tree of Life (38). The `RASE` reference database was constructed without additional ancestral state reconstruction as all resistance genotypes were known.

ST93 has two distinct clades, an ancestral MSSA clade with isolates from the Northern Territory, and a divergent MRSA clade, which expanded on the Australian East Coast, and spread to FNQ and PNG. This allowed us to assess genotyping ability of clade-specific methicillin resistance, which we have shown was compromised in the outbreak surveillance assessment using the approximate genomic neighbor typing approach in `Sketchy`. We expected `RASE` to have superior performance due to using a lineage phylogeny as guide for its genomic neighbor typing implementation with `ProPhyle 0.3.3.1` (28). To visualize the differences in resolution between our MinHash approach and tree-guided (SNP based) genomic neighbor typing (Fig. S4), we used `NetView` (39) to reconstruct genome population networks based on pairwise-distances from underlying SNPs and pairwise shared hash distance (*s* - *h*) computed with `Sketchy`. A value of *k* = 20 was selected for visualization of the network topologies in Fig. 5, as described previously (40) indicating stable configurations in both networks across selected community clustering algorithms (Fig. S1, C-D).

For comparison with `Sketchy`, we used a `RASE` (commit 27113cb) database constructed at *k* = 16, and the `Sketchy` outbreak reference sketch at *k* = 16 and *s* = 1000, as well as a high resolution sketch at *s* = 1000000. `RASE` requires sequence times per read, which were not available in the output of `Bonito v0.3.6`. We therefore used reads base called with the `Guppy v4.6` R9.4.1 DNA HAC model for this comparison. `RASE` outputs predictions by minute timestamps (including the number of reads) from which we selected the prediction closest to the 1000 read threshold (Table 7) used throughout this manuscript; we also ran the full read set to check for persistence of tetracycline prediction errors (Table S7). `RASE` read thresholds for each isolate were used for the read limit parameter (`--limit`) in the `Sketchy` predictions (Tables 8,9).

## ACKNOWLEDGMENTS

ES was funded by Queensland Genomics (41) and a joint PRISM2 & HOT North (NHMRC 1131932) pilot grant (42); ES, CH, EM were funded by a SERTA grant from Townsville University Hospital (2018_21); LC was supported by an NHMRC grant (NHMRC GNT1195743); PH, IA, AG, CF, RF, SS, ES, LC, SYCT, EM, WP were funded by a National Health and Medical Research Council Ideas grant (NHMRC 2012286)

---

**Fig. S1.**
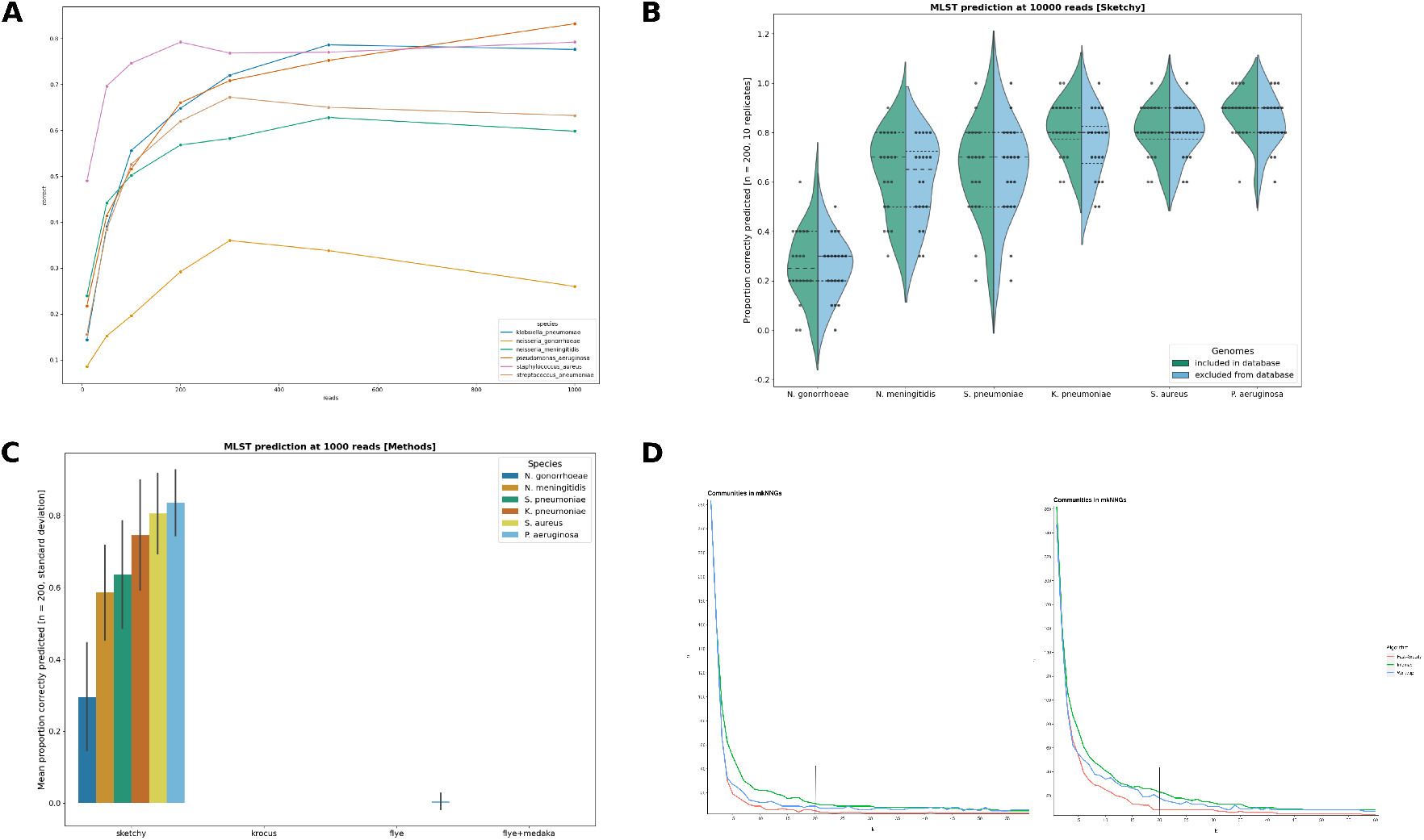
Supplementary figures for `Sketchy`: **(A)** MLST cross-validation prediction mean accuracy over a range of read thresholds (10 - 1000 reads, Methods) across the sic species outlined in Table 1, **(B)** MLST cross-validation prediction accuracy at 10000 reads across the six species, showing simulated nanopore runs when hold-out isolates were included (green) or excluded (blue) from the reference sketch, **(C)** MLST cross-validation prediction mean accuracy at 1000 reads (with standard deviation error bars) for `Sketchy` when compared to `Krocus` and typing from `Flye` and `Flye+Medaka` assemblies, **(D)** Mutual k-nearest-neighbor community assemblage plots using three different community detection algorithms over a range of k = 1 - 60 (left: shared hashes distance, right: SNP distance) indicating stable network topologies (Fig. 5) at the selected value (vertical lines).

## Data availability

Sequence data (Illumina, ONT) has been deposited under Bio-Project: PRJNA657380. Sketchy is open-source and available at: https://github.com/esteinig/sketchy. Reference genotypes and sequence data summaries can be found at the project repository: https://github.com/esteinig/ca-mrsa

## Supplementary Materials

**Table S1.**
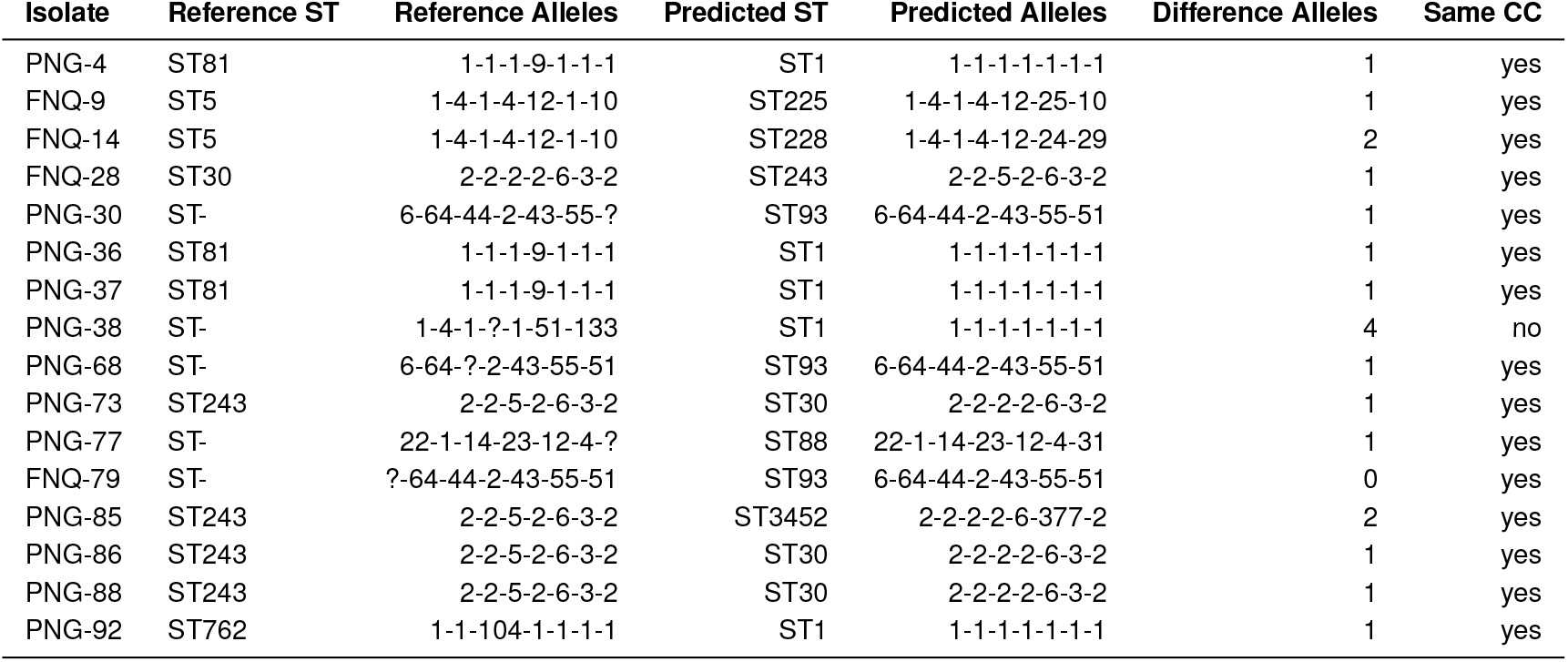
MLST error analysis from outbreak data predictions

| Isolate | Reference ST | Reference Alleles | Predicted ST | Predicted Alleles | Difference Alleles | Same CC |
| --- | --- | --- | --- | --- | --- | --- |
| PNG-4 | ST81 | 1-1-1-9-1-1-1 | ST1 | 1-1-1-1-1-1-1 | 1 | yes |
| FNQ-9 | ST5 | 1-4-1-4-12-1-10 | ST225 | 1-4-1-4-12-25-10 | 1 | yes |
| FNQ-14 | ST5 | 1-4-1-4-12-1-10 | ST228 | 1-4-1-4-12-24-29 | 2 | yes |
| FNQ-28 | ST30 | 2-2-2-2-6-3-2 | ST243 | 2-2-5-2-6-3-2 | 1 | yes |
| PNG-30 | ST- | 6-64-44-2-43-55-? | ST93 | 6-64-44-2-43-55-51 | 1 | yes |
| PNG-36 | ST81 | 1-1-1-9-1-1-1 | ST1 | 1-1-1-1-1-1-1 | 1 | yes |
| PNG-37 | ST81 | 1-1-1-9-1-1-1 | ST1 | 1-1-1-1-1-1-1 | 1 | yes |
| PNG-38 | ST- | 1-4-1-?-1-51-133 | ST1 | 1-1-1-1-1-1-1 | 4 | no |
| PNG-68 | ST- | 6-64-?-2-43-55-51 | ST93 | 6-64-44-2-43-55-51 | 1 | yes |
| PNG-73 | ST243 | 2-2-5-2-6-3-2 | ST30 | 2-2-2-2-6-3-2 | 1 | yes |
| PNG-77 | ST- | 22-1-14-23-12-4-? | ST88 | 22-1-14-23-12-4-31 | 1 | yes |
| FNQ-79 | ST- | ?-64-44-2-43-55-51 | ST93 | 6-64-44-2-43-55-51 | 0 | yes |
| PNG-85 | ST243 | 2-2-5-2-6-3-2 | ST3452 | 2-2-2-2-6-377-2 | 2 | yes |
| PNG-86 | ST243 | 2-2-5-2-6-3-2 | ST30 | 2-2-2-2-6-3-2 | 1 | yes |
| PNG-88 | ST243 | 2-2-5-2-6-3-2 | ST30 | 2-2-2-2-6-3-2 | 1 | yes |
| PNG-92 | ST762 | 1-1-104-1-1-1-1 | ST1 | 1-1-1-1-1-1-1 | 1 | yes |

**Table S2.** *S. aureus* outbreak isolates [streaming] (4 flow cells, 2 x 12-plex, 1000 reads, k = 16, s = 1000, n = 160)

| Feature | Binary | TP | TN | FP | FN | Accuracy (%) | Precision (%) | Sensitivity (%) | Specificity (%) |
| --- | --- | --- | --- | --- | --- | --- | --- | --- | --- |
| MLST | false | - | - | - | - | 87.50 | 87.91 | 87.50 | - |
| SCCmec type | false | - | - | - | - | 75.00 | 84.31 | 75.00 | - |
| PVL | true | 128 | 16 | 2 | 14 | 90.00 | 98.46 | 90.14 | 88.88 |
| Clindamycin | true | 1 | 141 | 12 | 6 | 88.75 | 7.69 | 14.28 | 92.15 |
| Rifampicin | true | 0 | 158 | 2 | 0 | 98.75 | na | na | 98.75 |
| Ciprofloxacin | true | 0 | 151 | 8 | 1 | 94.37 | na | na | 94.96 |
| Vancomycin | true | 0 | 160 | 0 | 0 | 100.0 | na | na | 100.0 |
| Tetracycline | true | 0 | 153 | 8 | 1 | 94.37 | na | na | 94.96 |
| Mupirocin | true | 0 | 157 | 1 | 2 | 98.12 | na | na | 99.36 |
| Gentamicin | true | 0 | 160 | 0 | 0 | 98.75 | na | na | 98.75 |
| Trimethoprim | true | 0 | 154 | 1 | 5 | 96.25 | na | na | 99.35 |
| Penicillin | true | 148 | 3 | 4 | 5 | 94.37 | 97.36 | 96.73 | 42.85 |
| Methicillin | true | 105 | 19 | 7 | 29 | 77.5 | 93.75 | 78.35 | 73.07 |
| Erythromycin | true | 1 | 141 | 12 | 6 | 88.75 | 7.69 | 14.28 | 92.15 |
| Fusidic Acid | true | 2 | 152 | 6 | 0 | 96.25 | 25.00 | 100.0 | 96.20 |

**Table S3.** *S. aureus* outbreak isolates on MinION (4 flow cells, 2 x 12-plex, 1000 reads, k = 16, s = 10000, n = 160)

| Feature | Binary | TP | TN | FP | FN | Accuracy (%) | Precision (%) | Sensitivity (%) | Specificity (%) |
| --- | --- | --- | --- | --- | --- | --- | --- | --- | --- |
| MLST | false | - | - | - | - | 93.75 | 90.53 | 93.75 | - |
| SCCmec type | false | - | - | - | - | 76.87 | 86.40 | 76.87 | - |
| PVL | true | 135 | 16 | 2 | 7 | 94.37 | 98.54 | 95.07 | 88.88 |
| Clindamycin | true | 0 | 149 | 4 | 7 | 93.12 | na | na | 97.38 |
| Rifampicin | true | 0 | 160 | 0 | 0 | 100.0 | na | na | 100.0 |
| Ciprofloxacin | true | 0 | 158 | 1 | 1 | 98.75 | na | na | 99.37 |
| Vancomycin | true | 0 | 160 | 0 | 0 | 100.0 | na | na | 100.0 |
| Tetracycline | true | 0 | 157 | 2 | 1 | 99.13 | na | na | 98.74 |
| Mupirocin | true | 0 | 158 | 0 | 2 | 98.75 | na | na | 100.0 |
| Gentamicin | true | 0 | 160 | 0 | 0 | 100.0 | na | na | 100.0 |
| Trimethoprim | true | 0 | 153 | 2 | 5 | 95.62 | na | na | 98.70 |
| Penicillin | true | 150 | 7 | 0 | 3 | 98.12 | 100.0 | 98.03 | 100.0 |
| Methicillin | true | 105 | 23 | 3 | 29 | 80.00 | 97.22 | 78.35 | 88.46 |
| Erythromycin | true | 1 | 149 | 4 | 6 | 93.75 | 20.00 | 14.28 | 97.38 |
| Fusidic Acid | true | 2 | 158 | 0 | 0 | 100.0 | 100.0 | 100.0 | 100.0 |

**Table S4.**
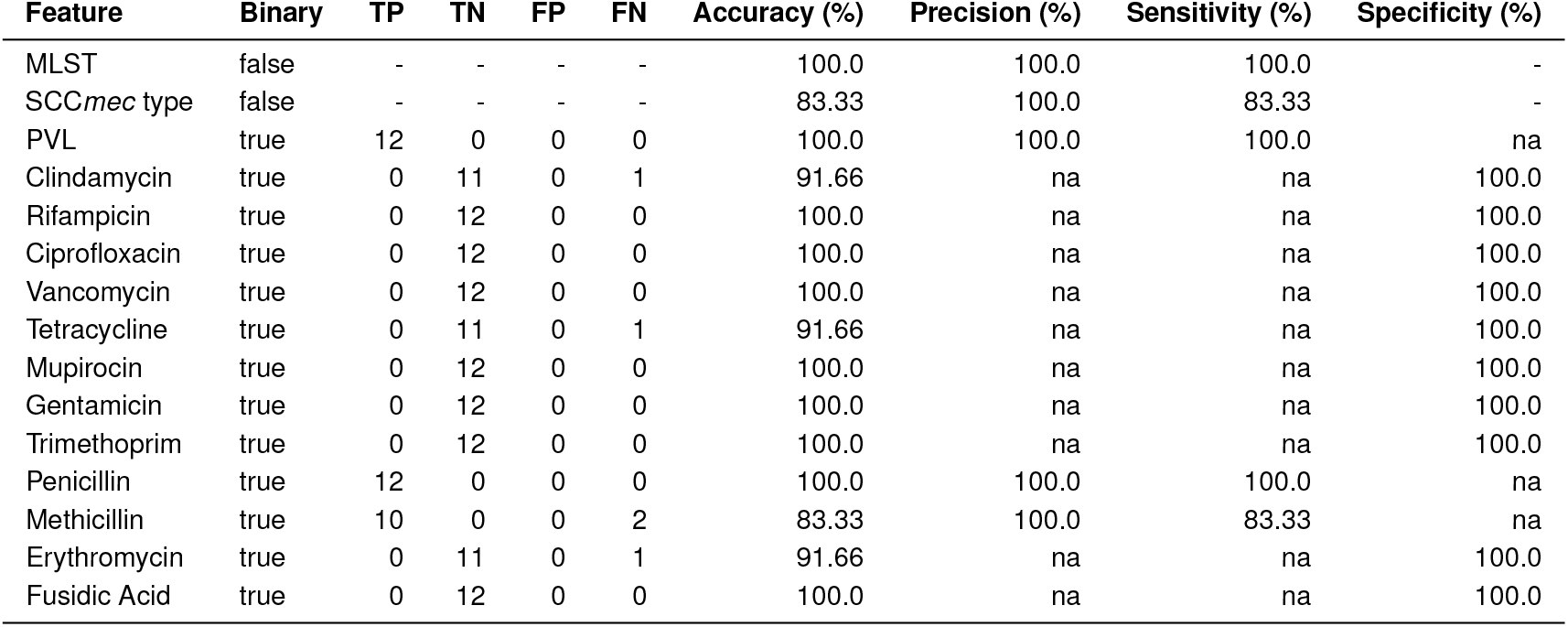
*S. aureus* outbreak isolates (Goroka) on MinION (1 flow cell, 12-plex, 1000 reads, k = 16, s = 10000, n = 12)

| Feature | Binary | TP | TN | FP | FN | Accuracy (%) | Precision (%) | Sensitivity (%) | Specificity (%) |
| --- | --- | --- | --- | --- | --- | --- | --- | --- | --- |
| MLST | false | - | - | - | - | 100.0 | 100.0 | 100.0 | - |
| SCCmec type | false | - | - | - | - | 83.33 | 100.0 | 83.33 | - |
| PVL | true | 12 | 0 | 0 | 0 | 100.0 | 100.0 | 100.0 | na |
| Clindamycin | true | 0 | 11 | 0 | 1 | 91.66 | na | na | 100.0 |
| Rifampicin | true | 0 | 12 | 0 | 0 | 100.0 | na | na | 100.0 |
| Ciprofloxacin | true | 0 | 12 | 0 | 0 | 100.0 | na | na | 100.0 |
| Vancomycin | true | 0 | 12 | 0 | 0 | 100.0 | na | na | 100.0 |
| Tetracycline | true | 0 | 11 | 0 | 1 | 91.66 | na | na | 100.0 |
| Mupirocin | true | 0 | 12 | 0 | 0 | 100.0 | na | na | 100.0 |
| Gentamicin | true | 0 | 12 | 0 | 0 | 100.0 | na | na | 100.0 |
| Trimethoprim | true | 0 | 12 | 0 | 0 | 100.0 | na | na | 100.0 |
| Penicillin | true | 12 | 0 | 0 | 0 | 100.0 | 100.0 | 100.0 | na |
| Methicillin | true | 10 | 0 | 0 | 2 | 83.33 | 100.0 | 83.33 | na |
| Erythromycin | true | 0 | 11 | 0 | 1 | 91.66 | na | na | 100.0 |
| Fusidic Acid | true | 0 | 12 | 0 | 0 | 100.0 | na | na | 100.0 |

**Table S5.** *S. aureus* outbreak isolates on Flongle (1 flow cell, 12-plex, 1000 reads, k = 16, s = 10000, n = 12)

| Feature | Binary | TP | TN | FP | FN | Accuracy (%) | Precision (%) | Sensitivity (%) | Specificity (%) |
| --- | --- | --- | --- | --- | --- | --- | --- | --- | --- |
| MLST | false | - | - | - | - | 100.0 | 100.0 | 100.0 | - |
| SCCmec type | false | - | - | - | - | 83.33 | 100.0 | 83.33 | - |
| PVL | true | 12 | 0 | 0 | 0 | 100.0 | 100.0 | 100.0 | na |
| Clindamycin | true | 0 | 11 | 1 | 0 | 91.66 | na | na | 91.66 |
| Rifampicin | true | 0 | 12 | 0 | 0 | 100.0 | na | na | 100.0 |
| Ciprofloxacin | true | 0 | 12 | 0 | 0 | 100.0 | na | na | 100.0 |
| Vancomycin | true | 0 | 12 | 0 | 0 | 100.0 | na | na | 100.0 |
| Tetracycline | true | 0 | 12 | 0 | 0 | 100.0 | na | na | 100.0 |
| Mupirocin | true | 0 | 12 | 0 | 0 | 100.0 | na | na | 100.0 |
| Gentamicin | true | 0 | 12 | 0 | 0 | 100.0 | na | na | 100.0 |
| Trimethoprim | true | 0 | 12 | 0 | 0 | 100.0 | na | na | 100.0 |
| Penicillin | true | 12 | 0 | 0 | 0 | 100.0 | 100.0 | 100.0 | na |
| Methicillin | true | 10 | 0 | 0 | 2 | 83.33 | 100.0 | 83.33 | na |
| Erythromycin | true | 0 | 11 | 1 | 0 | 91.66 | na | na | 91.66 |
| Fusidic Acid | true | 0 | 12 | 0 | 0 | 100.0 | na | na | 100.0 |

**Table S6.** Successive library experiment on MinION (1 flow cell, 4 x 12-plex, 1000 reads, k = 16, s = 10000, n = 48)

| Feature | Binary | TP | TN | FP | FN | Accuracy (%) | Precision (%) | Sensitivity (%) | Specificity (%) |
| --- | --- | --- | --- | --- | --- | --- | --- | --- | --- |
| MLST | false | - | - | - | - | 95.83 | 93.79 | 95.83 | - |
| SCCmec type | false | - | - | - | - | 77.08 | 93.62 | 77.08 | - |
| PVL | true | 46 | 0 | 1 | 1 | 95.83 | 97.87 | 97.87 | na |
| Clindamycin | true | 0 | 47 | 0 | 1 | 97.91 | na | na | 100.0 |
| Rifampicin | true | 0 | 48 | 0 | 0 | 100.0 | na | na | 100.0 |
| Ciprofloxacin | true | 0 | 47 | 0 | 1 | 97.91 | na | na | 100.0 |
| Vancomycin | true | 0 | 48 | 0 | 0 | 100.0 | na | na | 100.0 |
| Tetracycline | true | 0 | 47 | 0 | 1 | 97.91 | na | na | 100.0 |
| Mupirocin | true | 0 | 48 | 0 | 0 | 100.0 | na | na | 100.0 |
| Gentamicin | true | 0 | 48 | 0 | 0 | 100.0 | na | na | 100.0 |
| Trimethoprim | true | 0 | 48 | 0 | 0 | 100.0 | na | na | 100.0 |
| Penicillin | true | 48 | 0 | 0 | 0 | 100.0 | 100.0 | 100.0 | na |
| Methicillin | true | 36 | 1 | 1 | 10 | 77.08 | 97.29 | 78.26 | 0.50 |
| Erythromycin | true | 0 | 47 | 0 | 1 | 97.91 | na | na | 100.0 |
| Fusidic Acid | true | 0 | 48 | 0 | 0 | 100.0 | na | na | 100.0 |

**Table S7.** `RASE` classification of ST93 outbreak isolates, all reads (n = 120, lineage database)

| Feature | Binary | TP | TN | FP | FN | Accuracy (%) | Precision (%) | Sensitivity (%) | Specificity (%) |
| --- | --- | --- | --- | --- | --- | --- | --- | --- | --- |
| Clindamycin | true | 0 | 117 | 0 | 3 | 97.50 | na | na | 100.0 |
| Rifampicin | true | 0 | 120 | 0 | 0 | 100.0 | na | na | 100.0 |
| Ciprofloxacin | true | 0 | 120 | 0 | 0 | 100.0 | na | na | 100.0 |
| Vancomycin | true | 0 | 120 | 0 | 0 | 100.0 | na | na | 100.0 |
| Tetracycline | true | 0 | 82 | 37 | 1 | 68.33 | na | na | 68.91 |
| Mupirocin | true | 0 | 120 | 0 | 0 | 95.76 | na | na | 95.76 |
| Gentamicin | true | 0 | 120 | 0 | 0 | 100.0 | na | na | 100.0 |
| Trimethoprim | true | 0 | 120 | 0 | 0 | 100.0 | na | na | 100.0 |
| Penicillin | true | 120 | 0 | 0 | 0 | 100.0 | 100.0 | 100.0 | na |
| Methicillin | true | 117 | 0 | 3 | 0 | 97.50 | 97.50 | 100.0 | na |
| Erythromycin | true | 0 | 117 | 0 | 3 | 97.50 | na | na | 100.0 |
| Fusidic Acid | true | 0 | 120 | 0 | 0 | 100.0 | na | na | 100.0 |

## Notes

There are no conflicts of interest by the authors;

### Competing Interest Statement

The authors have declared no competing interest.

## References

1. Deamer D, Akeson M, Branton D (2016) Three decades of nanopore sequencing. Nat. Biotechnol. 34(5):518–524.

2. Samarakoon H, et al. (2020) Genopo: a nanopore sequencing analysis toolkit for portable android devices. Communications Biology 3(1):538.

3. Ferreira FA, Helmersen K, Visnovska T, Jørgensen SB, Aamot HV (2021) Rapid nanopore-based DNA sequencing protocol of antibiotic-resistant bacteria for use in surveillance and outbreak investigation. Microb Genom 7(4).

4. Steinig E, et al. (2021) Phylodynamic modelling of bacterial outbreaks using nanopore sequencing. bioRxiv.

5. Leggett RM, Heavens D, Caccamo M, Clark MD, Davey RP (2015) NanoOK: multi-reference alignment analysis of nanopore sequencing data, quality and error profiles. Bioinformatics 32(1):142–144.

6. Moss EL, Maghini DG, Bhatt AS (2020) Complete, closed bacterial genomes from microbiomes using nanopore sequencing. Nat. Biotechnol. 38(6):701–707.

7. Cao MD, et al. (2017) Scaffolding and completing genome assemblies in real-time with nanopore sequencing. Nat. Commun. 8(1):14515.

8. Nguyen SH, Duarte TPS, Coin LJM, Cao MD (2017) Real-time demultiplexing Nanopore barcoded sequencing data with npBarcode. Bioinformatics 33(24):3988–3990.

9. Nguyen SH, Cao MD, Coin LJM (2021) Real-time resolution of short-read assembly graph using ONT long reads. PLoS Comput. Biol. 17(1):1–18.

10. Cao MD, et al. (2016) Streaming algorithms for identification pathogens and antibiotic resistance potential from real-time MinION™ sequencing. Gigascience 5(1).

11. Dilthey AT, Jain C, Koren S, Phillippy AM (2019) Strain-level metagenomic assignment and compositional estimation for long reads with MetaMaps. Nat. Commun. 10(1):3066.

12. Hunt M, et al. (2019) Antibiotic resistance prediction for mycobacterium tuberculosis from genome sequence data with mykrobe [version 1; peer review: 2 approved, 1 approved with reservations]. Wellcome Open Research 4(191).

13. Charalampous T, et al. (2019) Nanopore metagenomics enables rapid clinical diagnosis of bacterial lower respiratory infection. Nat. Biotechnol. 37(7):783–792.

14. Whelan FJ, et al. (2020) Culture-enriched metagenomic sequencing enables in-depth profiling of the cystic fibrosis lung microbiota. Nature Microbiology 5(2):379–390.

15. Leggett RM, et al. (2020) Rapid MinION profiling of preterm microbiota and antimicrobial-resistant pathogens. Nature Microbiology 5(3):430–442.

16. Brinda K, et al. (2020) Rapid inference of antibiotic resistance and susceptibility by genomic neighbour typing. Nature Microbiology 5(3):455–464.

17. Aglua I, et al. (2018) Methicillin-Resistant Staphylococcus Aureus in Melanesian Children with Haematogenous Osteomyelitis from the Central Highlands of Papua New Guinea. Int. J. Pediatr. 6(10):8361–8370.

18. Guthridge I, Smith S, Horne P, Hanson J (2019) Increasing prevalence of methicillin-resistant *Staphylococcus aureus* in remote Australian communities: implications for patients and clinicians. Pathogen 51(4):428–431.

19. Wozniak TM, et al. (2020) Geospatial epidemiology of *Staphylococcus aureus* in a tropical setting: an enabling digital surveillance platform. Sci. Rep. 10(1):13169.

20. Steinig E, et al. (2021) Phylodynamic modelling of bacterial outbreaks using nanopore sequencing. bioRxiv.

21. Broder AZ (1997) On the resemblance and containment of documents in Proceedings. Compression and Complexity of SEQUENCES 1997 (Cat. No.97TB100171). pp. 21–29.

22. Andrei Z, Broder I (2000) Filtering Near-Duplicate documents, COM’00: Proceedings of the 11th annual symposium on combinatorial pattern matching.

23. Ondov BD, et al. (2016) Mash: fast genome and metagenome distance estimation using MinHash. Genome Biol. 17(1):132.

24. Ondov BD, et al. (2019) Mash screen: high-throughput sequence containment estimation for genome discovery. Genome Biol. 20(1):232.

25. Blackwell GA, et al. (2021) Exploring bacterial diversity via a curated and searchable snapshot of archived DNA sequences. bioRxiv.

26. van Hal SJ, et al. (2018) Global Scale Dissemination of ST93: A Divergent *Staphylococcus aureus* Epidemic Lineage That Has Recently Emerged From Remote Northern Australia. Front. Microbiol. 9:1453.

27. Steinig E, et al. (2021) Phylodynamic signatures in the emergence of community-associated MRSA. bioRxiv.

28. Brinda K, Salikhov K, Pignotti S, Kucherov G (2017) ProPhyle: a phylogeny-based metagenomic classifier using the Burrows-Wheeler Transform.

29. Brinda K, et al. (2020) Rapid inference of antibiotic resistance and susceptibility by genomic neighbour typing. Nature Microbiology 5(3):455–464.

30. Loose M, Malla S, Stout M (2016) Real-time selective sequencing using nanopore technology. Nat. Methods 13(9):751–754.

31. Lipworth S, et al. (2020) Optimized use of oxford nanopore flowcells for hybrid assemblies. Microb Genom 6(11).

32. Steinig E, Coin L (2022) Nanoq: ultra-fast quality control for nanopore reads. J. Open Source Softw. 7(69):2991.

33. Bainomugisa A, et al. (2018) Multi-clonal evolution of multi-drug-resistant/extensively drugresistant Mycobacterium tuberculosis in a high-prevalence setting of Papua New Guinea for over three decades. Microbial genomics 4(2):e000147.

34. Bradley P, et al. (2016) Rapid antibiotic-resistance predictions from genome sequence data for *Staphylococcus aureus* and *Mycobacterium tuberculosis*. Nat. Commun. 7:11465.

35. Kaya H, et al. (2018) SCC*mec*Finder, a Web-Based Tool for Typing of Staphylococcal Cassette Chromosome *mec* in *Staphylococcus aureus* Using Whole-Genome Sequence Data. mSphere 3(1):e00612–17.

36. Jolley KA, Bray JE, Maiden MCJ (2018) Open-access bacterial population genomics: BIGSdb software, the PubMLST.org website and their applications [version 1; peer review: 2 approved]. Wellcome Open Research 3(124).

37. Kolmogorov M, Yuan J, Lin Y, Pevzner PA (2019) Assembly of long, error-prone reads using repeat graphs. Nat. Biotechnol. 37(5):540–546.

38. Letunic I, Bork P (2019) Interactive tree of life (iTOL) v4: recent updates and new developments. Nucleic Acids Res. 47(W1):W256–W259.

39. Steinig EJ, Neuditschko M, Khatkar MS, Raadsma HW, Zenger KR (2016) netview p: a network visualization tool to unravel complex population structure using genome-wide SNPs. Mol. Ecol. Resour. 16(1):216–227.

40. Steinig EJ, et al. (2019) Evolution and Global Transmission of a Multidrug-Resistant, Community-Associated Methicillin-Resistant *Staphylococcus aureus* Lineage from the Indian Subcontinent. MBio 10(6).

41. Vidgen ME, et al. (2021) Queensland genomics: an adaptive approach for integrating genomics into a public healthcare system. npj Genomic Medicine 6(1):71.

42. Williams K, Rung S, D’Antoine H, Currie BJ (2021) A cross-jurisdictional research collaboration aiming to improve health outcomes in the tropical north of australia. The Lancet Regional Health – Western Pacific 9.

